# Parcellation of the human amygdala using recurrence quantification analysis

**DOI:** 10.1101/2020.09.10.291351

**Authors:** Krzysztof Bielski, Sylwia Adamus, Emilia Kolada, Joanna Rączaszek-Leonardi, Iwona Szatkowska

## Abstract

Several previous attempts have been made to divide the human amygdala into smaller subregions based on the unique functional properties of the subregions. Although these attempts have provided valuable insight into the functional heterogeneity in this structure, the possibility that spatial patterns of functional characteristics can quickly change over time has been neglected in previous studies. In the present study, we explicitly account for the dynamic nature of amygdala activity. Our goal was not only to develop another parcellation method but also to augment existing methods with novel information about amygdala subdivisions. We performed state-specific amygdala parcellation using resting-state fMRI (rsfMRI) data and recurrence quantification analysis (RQA). RsfMRI data from 102 subjects were acquired with a 3T Trio Siemens scanner. We analyzed values of several RQA measures across all voxels in the amygdala and found two amygdala subdivisions, the ventrolateral (VL) and dorsomedial (DM) subdivisions, that differ with respect to one of the RQA measures, Shannon’s entropy of diagonal lines. Compared to the DM subdivision, the VL subdivision can be characterized by a higher value of entropy. The results suggest that VL activity is determined and influenced by more brain structures than is DM activity. To assess the biological validity of the obtained subdivisions, we compared them with histological atlases and currently available parcellations based on structural connectivity patterns (Anatomy Probability Maps) and cytoarchitectonic features (SPM Anatomy toolbox). Moreover, we examined their cortical and subcortical functional connectivity. The obtained results are similar to those previously reported on parcellation performed on the basis of structural connectivity patterns. Functional connectivity analysis revealed that the VL subdivision has strong connections to several cortical areas, whereas the DM subdivision is mainly connected to subcortical regions. This finding suggests that the VL subdivision corresponds to the basolateral subdivision of the amygdala (BLA), while the DM subdivision has some characteristics typical of the centromedial amygdala (CMA). The similarity in functional connectivity patterns between the VL subdivision and BLA, as well as between the DM subdivision and CMA, confirm the utility of our parcellation method. Overall, the study shows that parcellation based on BOLD signal dynamics is a powerful tool for identifying distinct functional systems within the amygdala. This tool might be useful for future research on functional brain organization.

**Highlights:**

- A new method for parcellation of the human amygdala was developed
- The ventrolateral and dorsomedial subdivisions of the amygdala were revealed
- The two subdivisions correspond to the anatomically defined regions of the amygdala
- The two subdivisions differ with respect to values of entropy
- A new parcellation method provides novel information about amygdala subdivisions

## INTRODUCTION

The amygdala plays a crucial role in several emotion-related functions, such as reinforcement learning (Murray, 2007; Wassum and Izquierdo, 2015), social behavior (Adolphs, 2010), and emotion regulation (Blair et al., 2007), in both animals and humans (LeDoux, 2007; Janak & Tye, 2015). Moreover, dysfunction of the amygdala is associated with a variety of psychiatric disorders, such as autism spectrum disorders (ASDs), anxiety disorders, and depression (Phillips et al., 2003; Phelps & LeDoux, 2005). It has been shown that the anatomic basis of the diverse functions of the amygdala is its complex internal structure. According to animal studies, the amygdala comprises a set of small nuclei located very close to each other (Swanson & Petrovich, 1998). It is very likely that distinct groups of nuclei, comprising subdivisions of the amygdala, process different kinds of information. Thus, identifying the role of different subdivisions of the amygdala in humans might help in understanding both healthy emotional functioning and mechanisms of psychiatric disorders. However, distinct subdivisions of the human amygdala have not been clearly identified. The roles of the amygdala subdivisions are difficult to examine in humans mainly because of the lack of a generally accepted method of parcellating this structure (for a review see Kolada et al. 2017, pp. 123-124). In effect, several research groups have attempted to develop a common method that divides the human amygdala into smaller subparts on the basis of their unique structural or functional properties.

Structural parcellations are typically based on the idea that a given subdivision should exhibit distinct cytoarchitectonics, topography and structural connectivity. Cytoarchitecture-based parcellations involve differentiating subparts of the brain structure on the basis of the quantity, density and localization of different types of neuronal cells. To the best of our knowledge, only one study has used a cytoarchitectonic approach to delineate small subdivisions of the amygdala (Amunts et al. 2005). Amunts and colleagues performed postmortem analysis of brain tissue and defined borders of three subareas of the amygdala: the laterobasal, centromedial and superficial areas. The results of this analysis are available in the SPM Anatomy toolbox for future MRI-based research (Eickhoff, et al., 2005). Topography-based parcellations rely on the manual delineation of subdivisions of brain structures on the basis of the locations of other brain areas, blood vessels, and cerebrospinal fluid spaces located nearby. At least two previous studies have used this type of parcellation scheme for the amygdala (Entis et al.,2012; Tyszka & Pauli, 2016). However, topography-based methods are researcher-dependent and may lead to inconsistent results. Finally, parcellations based on structural connectivity are grounded in the assumption that different subdivisions of the brain areas correspond to different parts of brain circuits; thus, they might be characterized by different patterns of connectivity. In human research, it is possible to conduct structural connectivity analysis using diffusion tensor imaging (DTI). Regarding the amygdala, there are several ways to apply structural connectivity analysis (Bach et al.,2011; Solano-Castiella et al., 2010; Saygin et al., 2011). As connectivity-based methods largely rely on the way of estimating the connections between structures that has been selected, they yield distinct numbers and sizes of obtained parcels.

Functional parcellations are focused on the functional specialization of brain areas or the functional connectivity between them. If a given brain area is responsible for a specific function, it is possible to prepare a fMRI localization task to selectively activate this area. Although there is no specific task for the selective activation of distinct amygdala subdivisions, it is possible to perform meta-analysis of data derived from a large number of experiments, in which even low levels of activation of the amygdala have been detected. There are at least two works that have used this method for amygdala parcellation (Bzdok et al., 2013; Yang et al., 2016). Unfortunately, parcellations on the basis of functional specialization (especially meta-analysis) require a very large number of experiments. An alternative approach to the functional parcellation of the amygdala is parcellation based on its functional connectivity with other brain regions. Using resting-state fMRI (rsfMRI), which relies on the analysis of spontaneous fluctuations in BOLD signals in the absence of an explicit task, it is possible to identify functional connectivity between brain areas (i.e., functional interactions between them; Fox and Raichle 2007). To date, functional connectivity analysis has demonstrated that rsfMRI might be a valuable tool for dividing the amygdala into two (Mishra et al., 2014) or three parts (Bickart et al., 2014; Caparelli et al., 2017).

Although the above methods for functional parcellation have provided valuable insight into the functional heterogeneity in the human amygdala, they are encumbered by a common disadvantage: they have neglected the possibility that the spatial patterns of functional characteristics may quickly change over time. In recent years, however, there has been a growing number of neuroimaging studies that take into account the dynamics of changes in brain activity (Izhikevich et al., 2007). In fMRI studies, dynamics can be assessed by the sliding window approach in connectivity analysis (Hutchison, et al., 2013) or modeling with differential equations (Deco et al.,2008). This approach seems to be fruitful in the field of neuroimaging, and recently, it was used for parcellation of the human thalamus (van Oort et al., 2018; Kumar et al., 2017).

In the present study, we explicitly account for the dynamic nature of amygdala activity. We derive state-specific amygdala parcellations using recurrence quantification analysis (RQA), a nonlinear method derived from dynamic systems theory. Patterns of recurrence in a signal (understood as revisiting a previously visited state; Marwan & Webber, 2015) reflect the properties of underlying dynamics of a system, which generate the signal. RQA can be performed with short time series and allows us to assess multidimensional relations in signal features reflecting different aspects of recurrence. This method was successfully used for detecting task-related activity and for estimating functional connectivity patterns on the basis of fMRI data (Bianciardi, et al., 2008; Lombardi et al., 2017).

As animal and human research indicate that the amygdala consists of some subdivisions that differ in functional properties, we can expect that the dynamics of functional MRI signals differ by subdivision. It is possible to verify this idea by assessing rsfMRI data. During the resting state, different brain subsystems reveal relatively stable patterns of synchronous activity (De Luca et al., 2006; Damoiseaux et al., 2006). Thus, it might be possible to compute features of the dynamics of the BOLD signal from the resting state in all voxels within the amygdala and delineate borders between functionally distinct subdivisions. In the present work, we used rsfMRI data and RQA to estimate functionally distinct subdivisions of the amygdala based on the temporal characteristics of the signals. Analyses were performed on three datasets. To assess the biological validity of the obtained subdivisions, we compared these subdivisions with histological atlases and currently available parcellations (i.e., the anatomy probability map function based on structural connectivity and the SPM Anatomy toolbox). Moreover, we examined the cortical and subcortical projections of the subdivisions using accessible resting-state data. Our main goal was not only to develop another parcellation method but also to augment existing methods with novel information about the subdivisions. The results of the analysis on how patterns of activations within subdivisions change over time may reveal the dynamical organization of the functional subsystems that generate the signals.

## MATERIALS AND METHODS

### Subjects

In total, 102 subjects participated volunteered in the study and gave their informed consent on the documents prepared according to guidelines from the local ethics authorities from the University of Social Sciences and Humanities (SWPS University) in Warsaw and the University of Warsaw following guidelines of the Code of Ethics of the World Medical Association (Declaration of Helsinki). Individuals were recruited via an internet-based survey posted on social media. To participate in the study, the candidates had be aged 21 – 35 years and right-handed; those who had metal medical devices located in the body, claustrophobia, psychiatric or neurological impairments and tattoos or permanent make-up were excluded from the study.

1. <u>The first dataset</u> included data of 37 subjects (18 males, mean age: 25 (SD: 2,19), 19 females, mean age: 24,95 (SD: 3,33); general mean age: 24,97 (SD: 2,8)).
2. <u>The second dataset</u> included data of 35 subjects (17 males, mean age: 24,56 (SD: 3,22), 18 females, mean age: 24,11 (SD: 2,00); general mean age: 24,25 (SD: 2,67)).
3. <u>The third dataset</u> included data of 30 subjects (15 males, mean age: 25, (SD: 3,72), 15 females, mean age: 24,5 (SD: 2,71); general mean age: 25 (SD: 3,18)).

### Data

All datasets were acquired using a Siemens Trio 3T scanner equipped with a 32-channel head coil. The scan parameters were as follows:

1. <u>The structural scan parameters (all datasets)</u> included T1w images with an isotropic voxel size of 1 mm^3^ (TR = 2.53 s, TE = 3 ms, flip angle = 7 degrees, FOV =256 mm, imaging matrix size = 256 x 256 x 176 mm, time of acquisition: 6 min)
2. <u>The functional scan parameters (first and second datasets)</u> included a scan time of 15 minutes in the resting state with the participant’s eyes opened (fixating on the cross), an isotropic voxel size of 2.5 mm x 2.5 mm x 2.5 mm (TR = 1.5 s, TE = 29 ms, FOV = 192 x 192 mm, imaging matrix size = 96 x 96 x 80 mm), and 593 volumes.
3. <u>The functional scan parameters (third dataset)</u> included a scan time of 15 minutes in the resting state with the participant’s eyes opened (fixating on the cross), an isotropic voxel size of 3 mm x 3 mm x 3 mm (TR = 1,25 s, TE = 29 ms, FOV = 192 x 192 mm, imaging matrix size = 96 x 96 x 80 mm), and 586 volumes.

### Preprocessing

Preprocessing was performed with a default preprocessing pipeline for volume-based analyses using the CONN functional connectivity toolbox (Whitfield-Gabrieli, S., & Nieto-Castanon, A. 2012). First, the realign & unwarp procedure from SPM12 (Andersson et al., 2001) was run on the functional volumes. Next, the data were segmented into white matter, gray matter and cerebrospinal fluid and normalized into a standard MNI152 2 mm space. Finally, the data were spatially smoothed with an 8 mm full width at half maximum Gaussian kernel and denoised with a bandpass filter between 0.01 and 0.1 Hz to minimize the effects of physiological noise.

### Data analysis - pipeline

Figure 1 presents the general schema of the analysis. First, the parameters for the RQA were estimated to create the recurrence plots (RPs), and the RQA measures were computed on the signal from each voxel from all subjects included in the first dataset. Voxels within the amygdala were then grouped into clusters based on the RQA measures values (orange frames on Figure 1), and the solution with the best internal validity coefficients was selected. To confirm the results, the analysis was run on another dataset (second dataset) using the same parameters and previously selected RQA measures (blue frames). Finally, the solutions from both datasets were compared with those of existing parcellations, and we investigated the functional connectivity patterns of the obtained subdivisions (green frames) with the third dataset to validate the anatomical reliability of the obtained solution and obtain additional insight into the functional organization of the human amygdalae. In the next subsections of this paper, there are detailed descriptions of the stages.

**Fig. 1.**
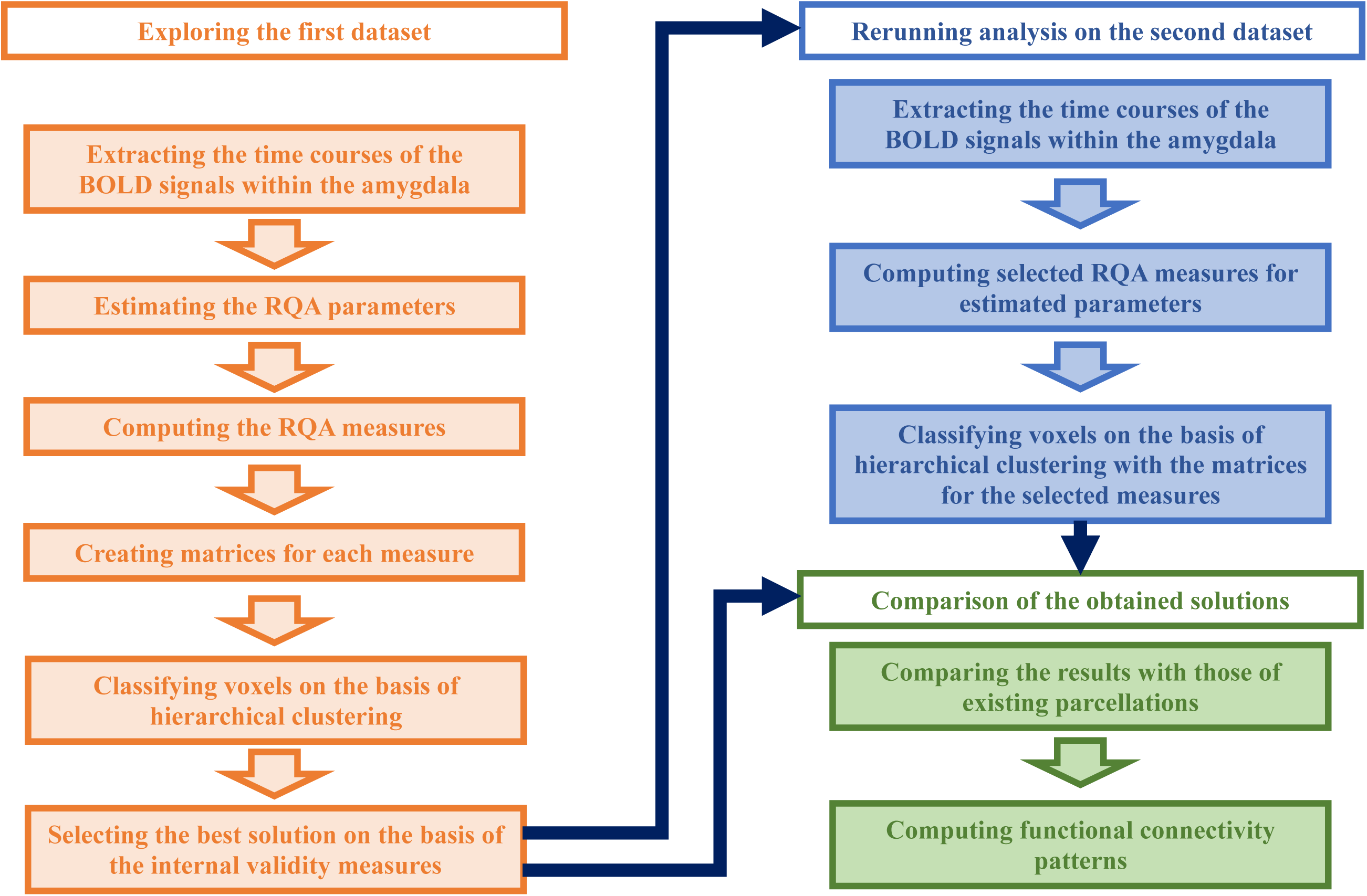
The pipeline for the whole analysis. The analysis was divided into three main stages: exploring the first dataset (orange frames), rerunning the analysis on the second dataset (blue frames), comparing the obtained solutions and computing functional connectivity patterns for the third dataset (green frames).

### Data analysis - masks

Voxels representing the amygdala were extracted from the structural and functional data by the anatomy probability map function based on the structural connectivity (APMC) mask prepared by Bach and collaborators (2011). The outline of this mask was drawn manually for each individual and then parcellated into two subdivisions on the basis of the structural connectivity patterns derived by diffusion weighted imaging (DWI). This mask was chosen because it allows the results obtained on the basis of functional features to be compared directly with those obtained by parcellation performed with structural data. Since the structural representation of the brain areas is not completely congruent with the functional representation, a comparison of these two perspectives might enrich our analysis.

### Data analysis - recurrence quantification analysis (RQA)

To assess the dynamical features of the BOLD signals, we applied recurrence quantification analysis (RQA) to all time series from the voxels of the amygdala defined, as described in the previous subsection. RQA focuses on characterizing the dynamics of a system with quantitative measures of recurrence. To calculate the values, it is necessary to recreate the trajectory of the system in a multidimensional phase space by using the acquired time series. Subsequently, on the basis of the trajectory in the multidimensional phase space, a recurrence plot (RP) is created. This plot presents the recurrence of particular states in the phase space trajectory as points on a two-dimensional space. Figure 2 presents an example of a recurrence plot. The RP was created and all computations in the RQA (except one stage of the analysis – look in the supplementary materials) were performed using the CRP toolbox (Marwan et al., 2007) operating under the MATLAB.

**Fig. 2.**
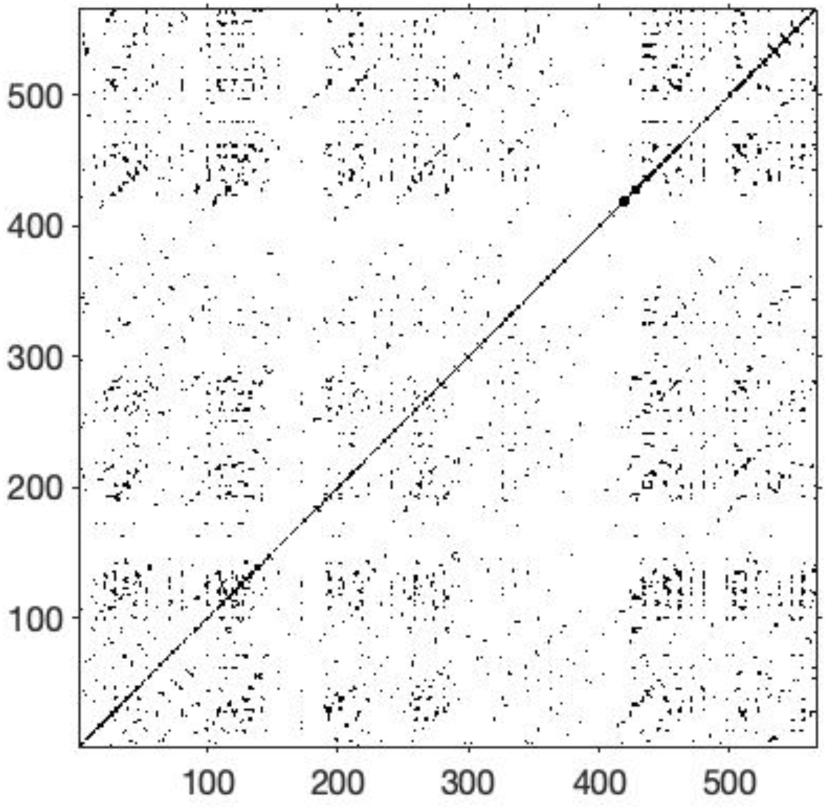
An example of a recurrence plot generated from randomly chosen time series from the first dataset.

Recurrence quantification refers to the quantification of particular structures on this plot, which are informative of the system’s dynamics. Details of the RQA are provided in the Supplementary Materials (SM.1.1. – Theoretical Background). We computed the following measures: determinism (%DET), the average diagonal line, the maximum length of the diagonal line, Shannon’s entropy of diagonal lines (ENTR), laminarity (LAM) and the trapping time (TT). The first four measures describe how predictable/unpredictable the trajectory of the system is in the phase space, while the last two measures provide information regarding one state for a given period of time. Details are provided in the Supplementary Materials (SM.1.3 – RQA measures). Each RQA measure was normalized across all voxels for each subject (mean value = 0, standard deviation = 1).

### Data analysis – software

The remaining stages of the analysis (except connectivity) were performed using Python 3.7 operating under Anaconda Jupyter Notebook 6.0.1 using packages such as: NumPy 1.17.2 (Oliphant, 2006), nibabel 3.0.0 (Brett et al., 2019), scikit-learn 0.21.3 (Pedregosa et al., 2011), scipy 1.3.1 (Virtanen et al., 2020) and matplotlib 3.0.3 (Hunter, 2007).

### Data analysis – Clustering

We designed six matrices, one for each measure (example shown in Figure 3). The rows represent the subjects, while the voxels in the MNI space correspond to consecutive columns. We assumed that the distribution of the RQA measures in the population could be a good indicator of these dynamics. As the dynamics of the signal in different subdivisions should be similar in the whole population, it might be possible to classify voxels into clusters representing functionally separate subdivisions on the basis of at least one RQA measure. To determine which measure is the best for this purpose, we tested all of them and selected the most appropriate one. For classification purposes, we used hierarchical clustering, which divides the dataset into smaller subparts on the basis of similarity. The similarity between vectors was assessed with ward metrics. Additionally, we imposed spatial constraints: two points could connect only if they were immediate neighbors to each other (voxels located next to each other), and the number of clusters was set to two.

**Fig. 3.**
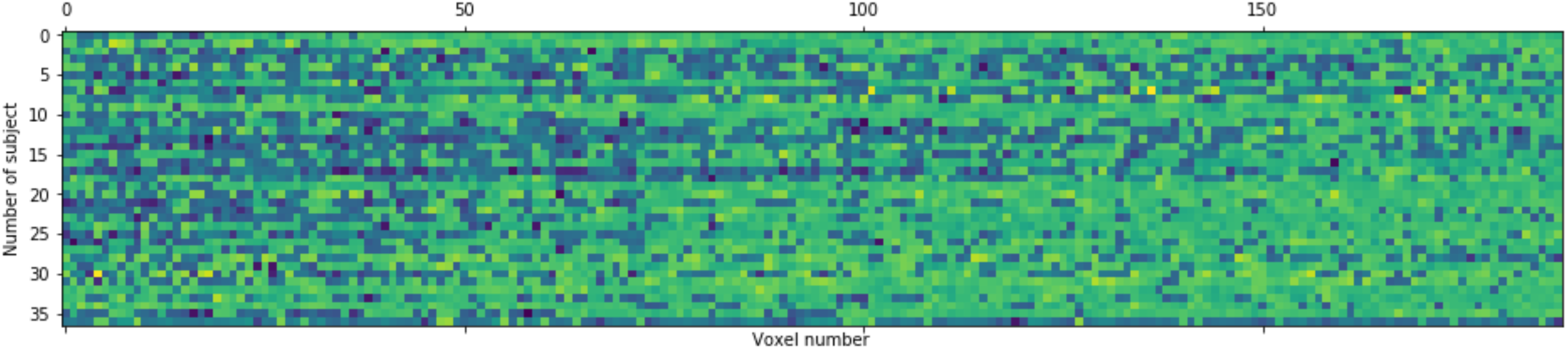
Example of a matrix used for clustering. The x axis corresponds to the voxels in the MNI space, and the y axis corresponds to the subjects in the analysis. The matrix was generated using the Matplotlib toolbox with Python 3.7.

### Data analysis - Selection of the solution - Internal Validity

To choose the most appropriate solution, we computed several metrics of internal validity for all 6 solutions explored in this paper. The internal validity metrics describe the location of voxels within and between clusters. The ideal solution is identified when voxels within a cluster are similar to each other and the voxels between clusters are very different. We used several measures of internal validation, such as the BetaCV, normalized cut and silhouette coefficient (for descriptions, see the Supplementary Materials – SM.2. Internal Validation Measures). However, we computed these measures using the distances in the spatial space with respect to the locations in the brain rather than the distances in the data space. In other words, the voxels were classified into different subdivisions by the clustering algorithm on the basis of the distances between vectors and values of the RQA measures among subjects in the given dataset. The more similar the values were between vectors, the more likely they were located within one cluster. On the other hand, we looked for the subdivisions that had compact spatial structures, not clusters with voxels distributed randomly across the amygdala. Thus, we decided to choose this parcellation, which had the most appropriate values of the internal validity measures for the distances in the brain space. In subsequent analyses, we used only these clustering solutions, which can be characterized by the highest (normalized cut, silhouette coefficient) or the lowest (BetaCV) values of the internal validation measures among the possible solutions derived from the first dataset.

### Data Analysis – performing analysis on the second dataset

To prove the reliability and stability of the amygdala parcellation, we repeated the analysis with the same parameters and the selected RQA measure on another dataset (the second dataset). Additionally, we compared our results with those of existing parcellations and performed connectivity analysis to determine the functional connectivity patterns of different subdivisions (using the third dataset).

### Data Analysis - comparison with existing parcellations

To assess the anatomical validity of the obtained solutions, we visualized our results and determined whether the obtained parts were classified into the same subdivision in the left and right amygdalae (we computed proportions of the given solutions and measures of similarity taken from the external validation measures – SM.3.). Furthermore, we compared our results obtained on the basis of the similarity of vectors regarding the distribution of the RQA measures across populations with those of existing parcellations of the amygdala: parcellation on the basis of cytoarchitectonic features provided in the SPM Anatomy toolbox (Amunts et al., 2005) and parcellation on the basis of structural connectivity patterns assessed via diffusion weighted imaging (Bach et al., 2011). Comparisons were made using indices such as purity, mutual information, the Jaccard coefficient and the Fowlkes–Mallows score (for details, see the Supplementary Materials – SM.3. External Validation Measures).

### Data Analysis - Connectivity Analysis

To determine whether the resulting subdivisions of the amygdala corresponded to anatomical groups of nuclei, which can be characterized by distinct connectivity patterns, we performed ROI-to-ROI functional connectivity analysis on the third dataset using the common part of the obtained subdivisions in the first and second datasets as masks of the different amygdala subparts. Our analysis was conducted with the CONN toolbox, and we used the Harvard-Oxford Atlas for the cortical and subcortical regions.

## RESULTS

### Exploration of the first dataset

#### Selection of the solution on the basis of internal validity

Based on the values of the internal validity coefficients, the parcellation solutions were ranked (the full tables with rankings are provided in the Supplementary Materials – SM.2.). The best voxel classification for all coefficient rankings was that based on Shannon’s entropy of diagonal lines (the exact values of the internal validity coefficients for this solution are provided in Table 1). The BetaCV measure for the best solution showed that the general intercluster distances are approximately 61% larger than the intracluster distances. Similar findings were observed when the values of the normalized cut measure and the most restrictive indicator – the silhouette coefficient (SC)

**Table 1.** Values of the internal validation measures for the selected solution for the first dataset.

| internal validation measures | left amygdala | right amygdala |
| --- | --- | --- |
| BetaCV | 0.615287 | 0.605200 |
| normalized cut | 0.766615 | 0.766903 |
| silhouette coefficient | 0.355453 | 0.359694 |

– were assessed. Additionally, the SC being equal to approximately 0.36 suggests that our results are not random.

#### Solution obtained on the first dataset

The resulting parcellation of the amygdala, as presented in Figure 4, divides the structure into two subdivisions. The ventrolateral subdivision (VL, green subdivision in the figure) includes 95 voxels from the left amygdala and 87 voxels from the right amygdala, and the dorsomedial (DM) subdivision consists of 93 voxels from the left amygdala and 65 voxels from the right amygdala.

**Fig. 4.**
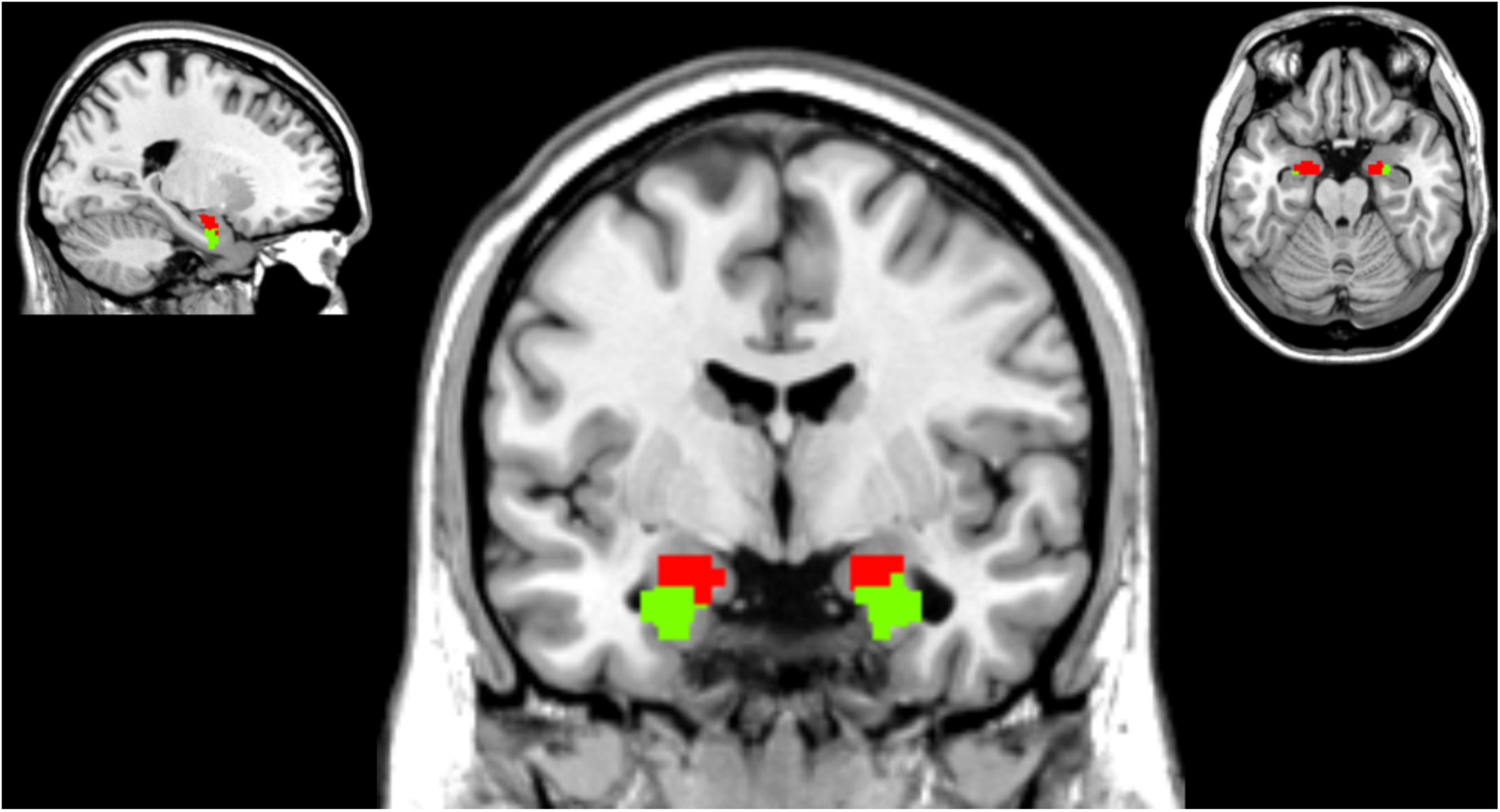
The results of our analysis were visualized by the amygdala mask reported in the publication by Dominik Bach and colleagues (2011) on the first dataset. Coordinates: x = 20, y = -4, and z = -20; views: sagittal, coronal, and axial.

The distribution of the entropy values in the amygdala voxels was left skewed (Figure 5). This pattern suggests that the values of entropy were higher in the VL part of the amygdala than in DM part of the amygdala (Figure 6).

**Fig. 5.**
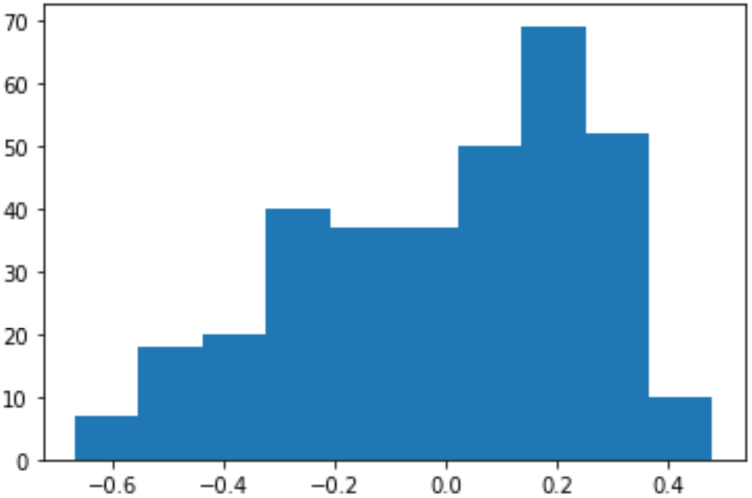
Histogram of the entropy values in voxels of the left and right amygdala subdivisions of the parcellation for the first dataset.

**Fig. 6.**
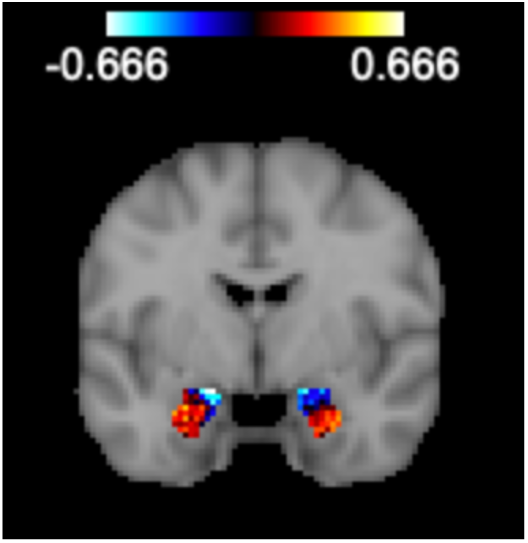
Distribution of the values of entropy in voxels of the left and right amygdala subdivisions of the parcellation for the first dataset.

#### Differences in measures between the two subregions

The mean values of entropy for each subdivision are presented in Table 2. The VL subdivision can be characterized by positive values of entropy, while the DM subdivision usually has negative values for the normalized entropy (Figure 6). Positive values in the distribution mean that the value of entropy is higher than the average value of entropy in the amygdala, while voxels with negative values have entropy values that are smaller than the average entropy value in a subject. These results were statistically significant.

**Table 2.** Mean values of entropy in the amygdala subdivisions obtained for the first dataset.

| amygdala | first cluster (ventrolateral) |  |  | second cluster (dorsomedial) |  |  |
| --- | --- | --- | --- | --- | --- | --- |
|  | left | right | both | left | right | both |
| mean entropy value | 0.1901 | 0.1630 | 0.1771 | -0.1941 | -0.2182 | -0.2041 |

The values of entropy significantly differed between the left VL and the left DM parts in the first dataset (Welch’s t test for independent samples, t= 16.21, p < 0.0001). Similarly, the values of entropy significantly differed between the right VL and the right DM subdivisions (t=11.4634, p<0.0001). However, the entropy levels between the left and right VL subdivisions (t =1.258, p =0.21) and between the left and right DM subdivisions did not significantly differ (t=0.7791, p-value=0.437). These results are shown in graphical form in Figure 7. All computations were performed using the SciPy toolbox from Python 3.7.

**Fig. 7.**
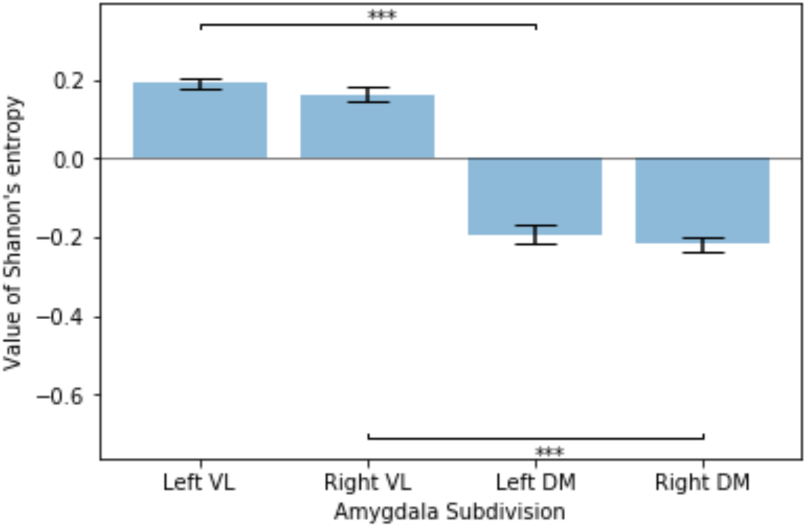
Mean values of entropy with standard errors for the amygdala subdivisions obtained for the first dataset.

### Repeated Analysis on the Second Dataset

To verify the stability of the solution, the whole procedure was carried out on the second dataset using the same RQA parameters, and Shannon’s entropy of diagonal lines was used to differentiate the amygdala subdivisions. The results of this confirmatory analysis are presented in Figure 8. Similar to the previously obtained results, the VL part of the amygdala (left: 128 voxels, right: 101 voxels) had a larger volume than did the DM part (left: 60 voxels, right: 51 voxels).

**Fig. 8.**
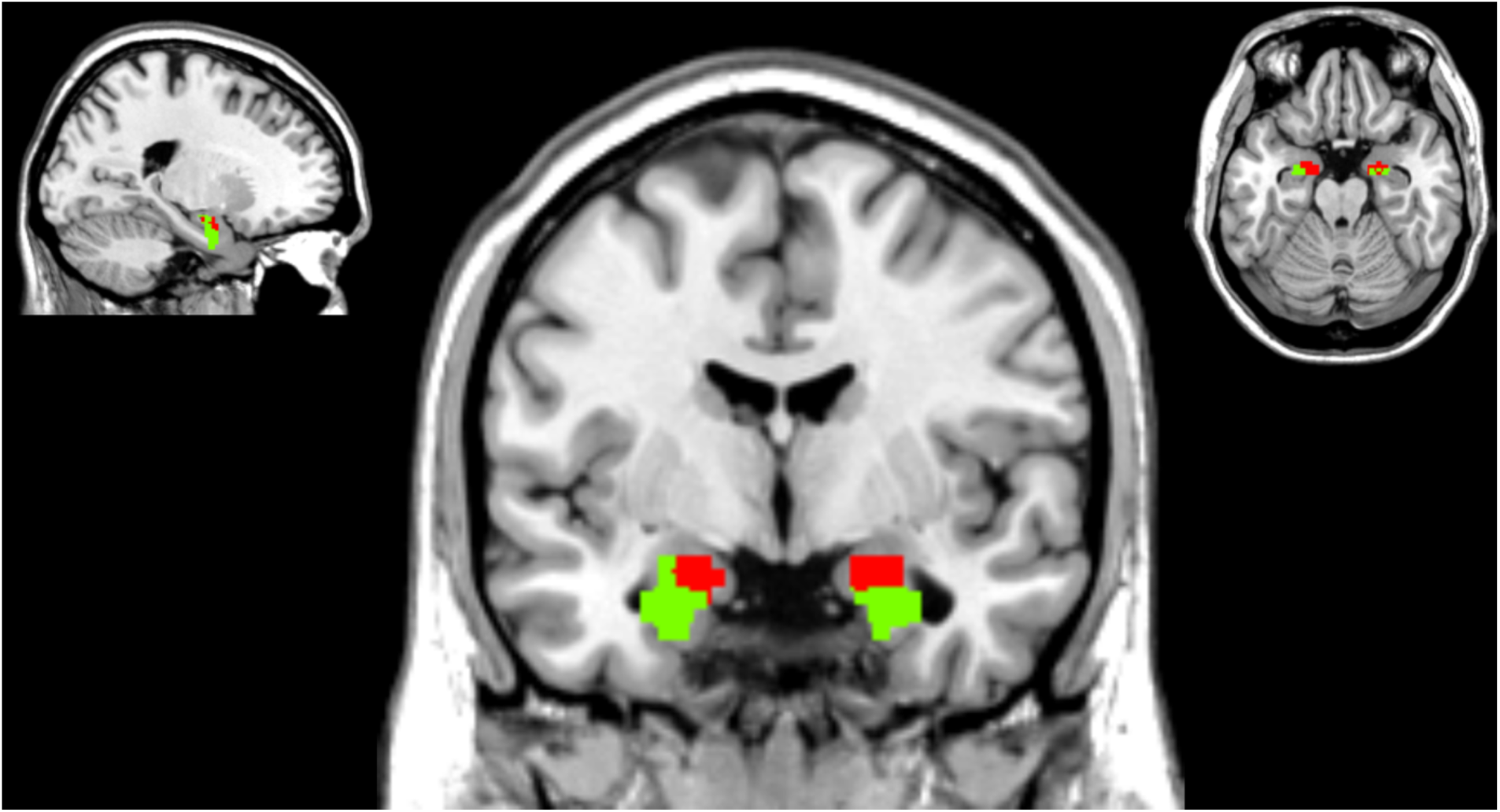
The results of our analysis were visualized by the amygdala mask reported in the publication by Dominik Bach and colleagues (2011) on the second dataset. Coordinates: x = 20, y = -4, and z = -20; views: sagittal, coronal, and axial.

The distribution of the entropy values in the amygdala voxels in the second dataset (Figure 9) had a similar tendency to be left-skewed, but in general, the values were smaller than those in the first dataset. For that reason, some voxels with negative values of entropy might have been classified by the algorithm as being in the VL subdivision (Figure 10). Thus, the spatial distributions of the entropy values for the two datasets seemed to be reasonably similar. The mean square error of the entropy values was smaller than 5%.

**Fig. 9.**
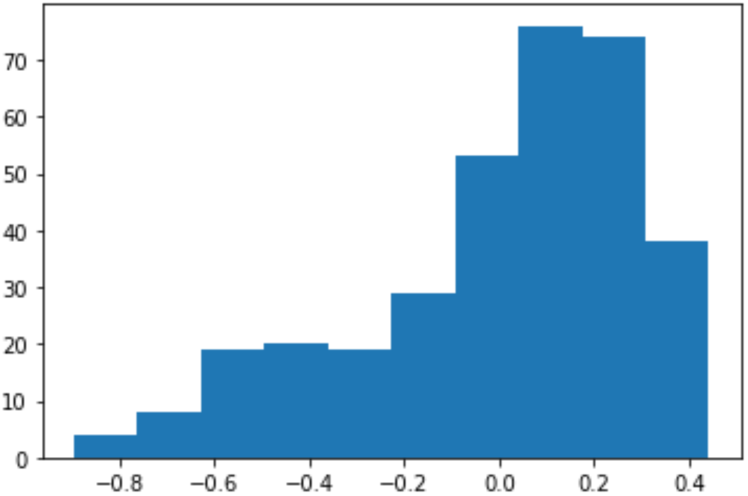
Histogram of the entropy values in voxels of the left and right amygdala subdivisions determined by parcellation for the second dataset.

**Fig. 10.**
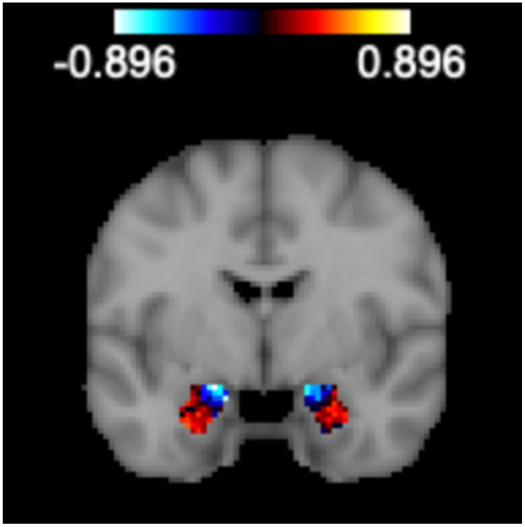
Distribution of the values of entropy in voxels of the left and right amygdala subdivisions determined by parcellation for the second dataset.

Figure 10 shows that the ventrolateral part can be characterized by positive values of entropy, while the DM part has predominantly negative values of entropy. These differences between the two subdivisions were statistically significant in both the left and right amygdalae (left amygdala: t = 16.1064, p <0.001; right amygdala: t=10.7475, p <0.001). There were no significant differences in the values of entropy between the left and right VL subdivisions (t=1.6335, p=0.104) or between the left and right medial inferior subdivisions (t= -1.9784, p=0.0505). These results are presented in graphical form in Figure 11.

**Fig. 11.**
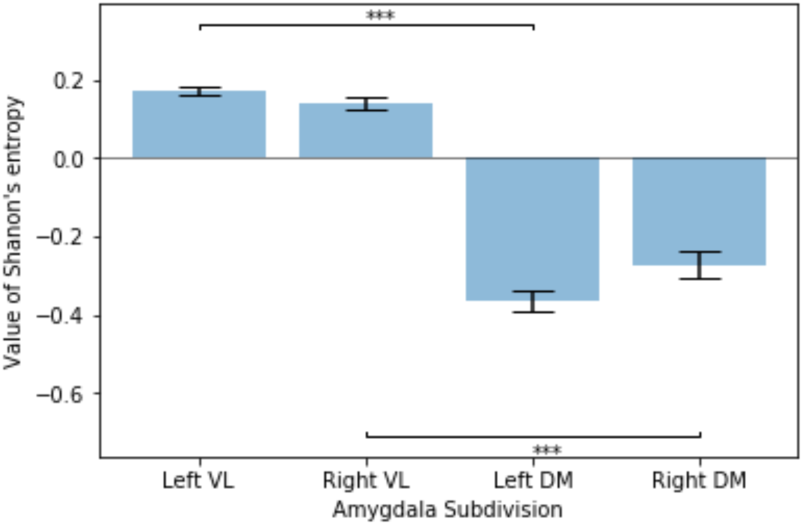
Mean values of entropy with standard errors for the amygdala subdivisions obtained for the second dataset.

### Comparisons of the Obtained Solutions

#### Spatial Representation

Here, we present a quantitative comparison of the parcellation solution for the first dataset, the parcellation solution for the second dataset and those of existing parcellations: structural connectivity-based parcellation (AMPC mask) by Bach et al. (2011) and the cytoarchitectonic atlas of the amygdala by Amunts et al. (2005) generated with the Anatomy toolbox (AT mask).

Additionally, we show the similarities between first and second dataset results (voxels that were classified to the same subdivisions in both analyses, see Figures 12 and 13).

**Fig. 12.**
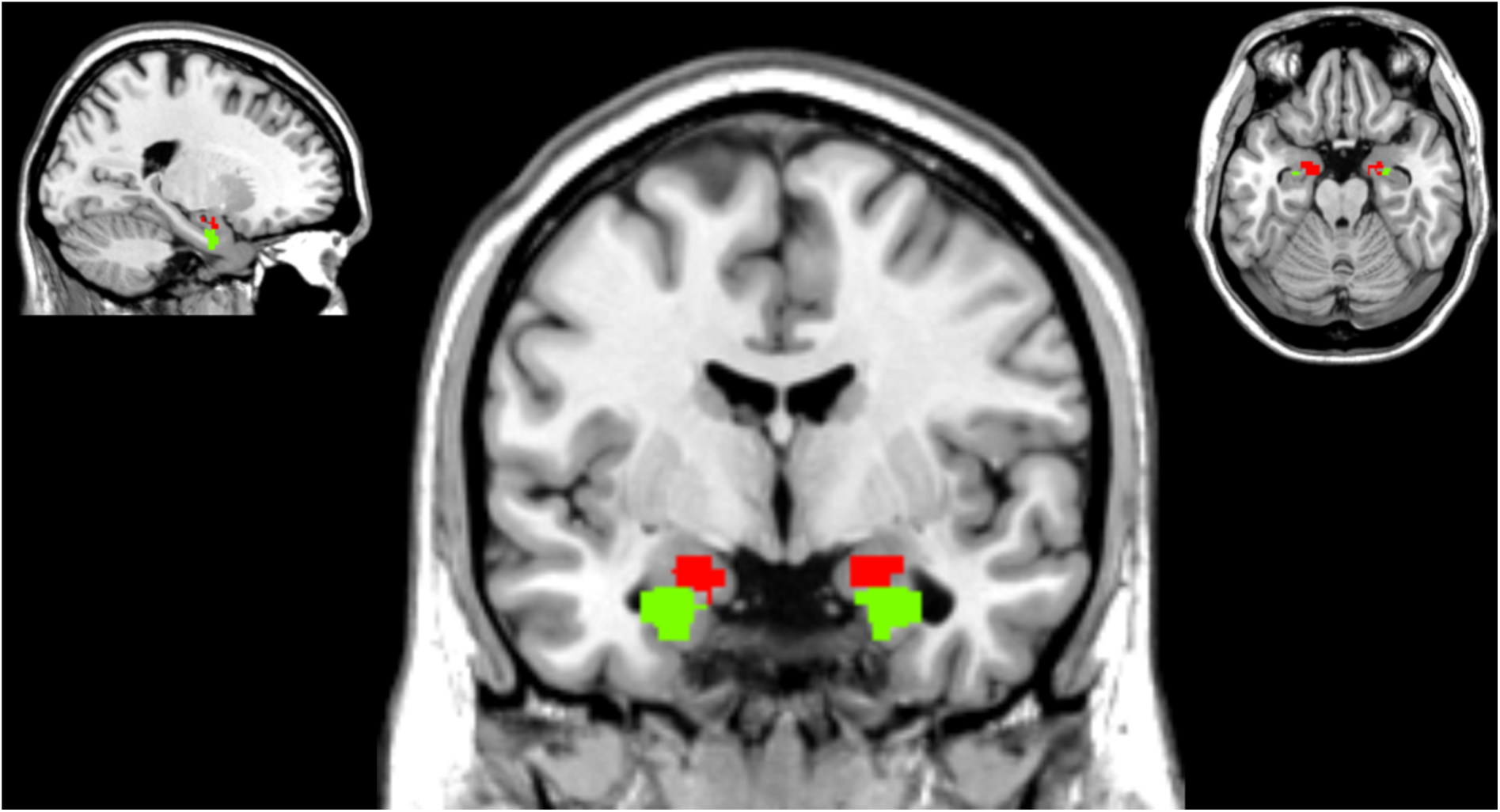
Visualization of the similarities between the first and second datasets in our results obtained with the amygdala mask reported in the publication by Dominik Bach and colleagues (2011). Coordinates: x = 20, y = -4, and z = -20; views: sagittal, coronal, and axial.

**Fig. 13.**
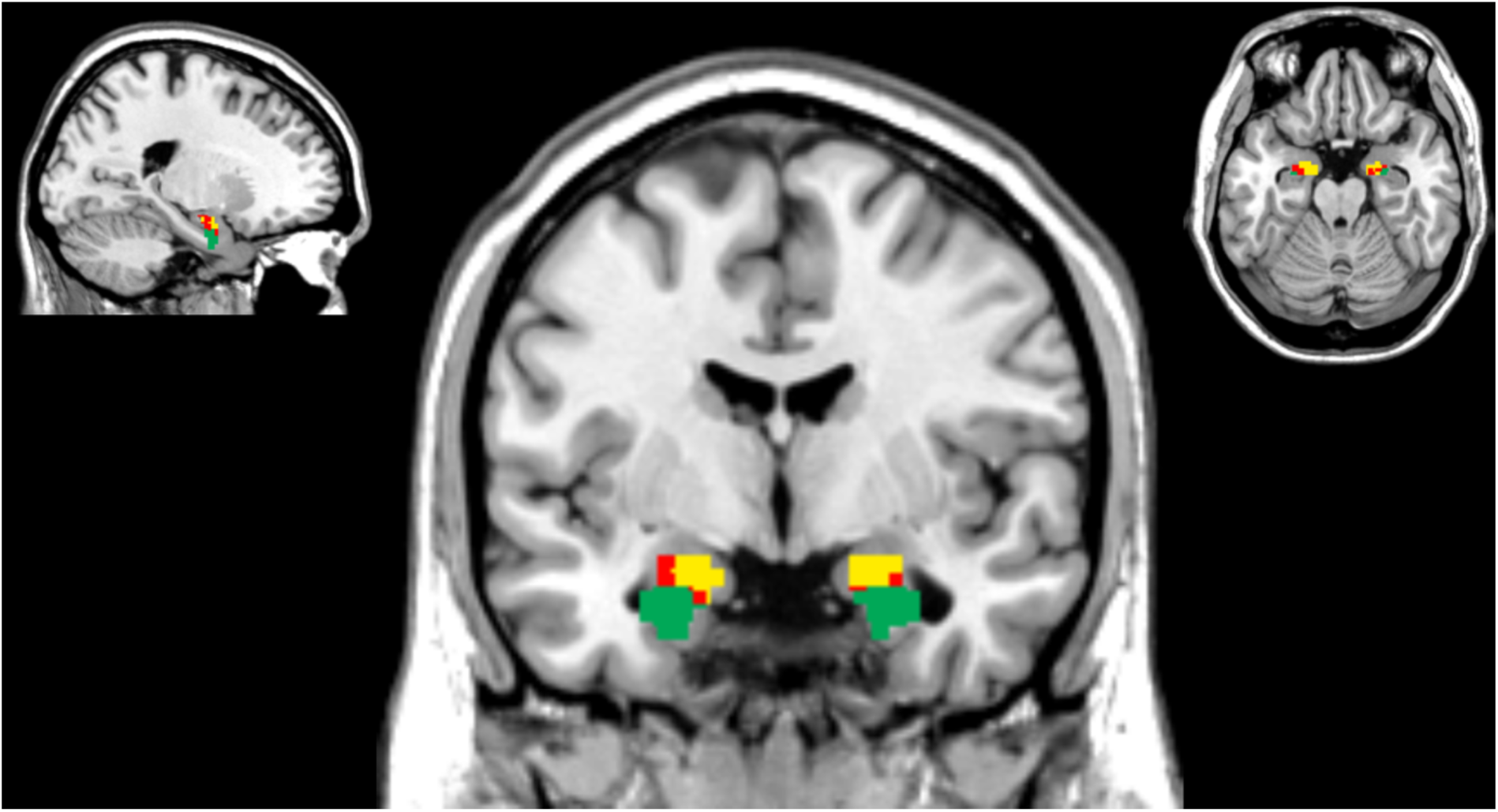
Visualization of the similarities between the first and second datasets in our results obtained with the amygdala mask reported in the publication by Dominik Bach and colleagues (2011), indicated by the green and yellow parts. The red regions differ between the two parcellations. Coordinates: x = 20, y = -4, and z = -20; views: sagittal, coronal, and axial.

The proportions of the sizes of the subdivisions obtained in the analysis of the first dataset correspond well with those of the deep and superficial subdivisions from the structural connectivity patterns (Bach et al.,2011). On the other hand, the results obtained for the second dataset show similar proportions of sizes to those of the laterobasal and joint centromedial and superficial subdivisions in the solution presented by Amunts and colleagues (2005) (Table 4).

**Table 3.**
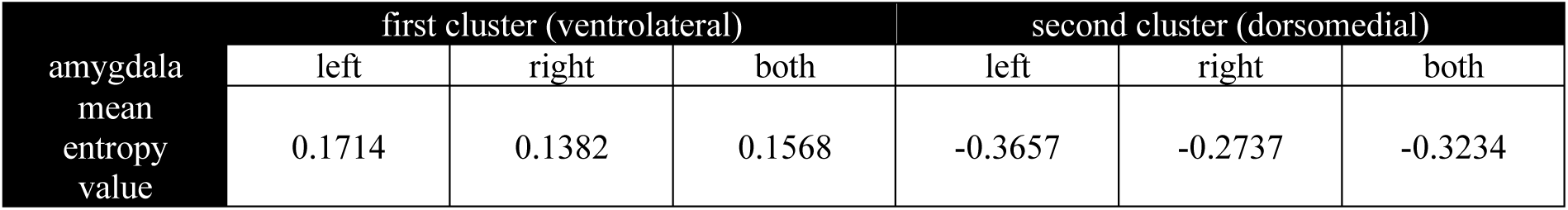
Mean values of entropy in the amygdala subdivisions obtained for the second dataset.

**Table 4.** Proportions of the sizes of different amygdala subregions across parcellations for the first and second datasets and those obtained by Bach et al. (2011) and Amunts et al. (2005).

|  | First dataset |  | Second dataset |  | Common Ground |  | Bach et al. (2011) |  | Amunts et al. (2005) |  |
| --- | --- | --- | --- | --- | --- | --- | --- | --- | --- | --- |
|  | VL | DM | VL | DM | VL | DM | Deep | Superficial | Laterobasal | Centromedial + Superficial |
| Left | 0.5053 | 0.4947 | 0.6809 | 0.3191 | 0.6206 | 0.3793 | 0.5160 | 0.4840 | 0.7647 | 0.2353 |
| Right | 0.5724 | 0.4276 | 0.6645 | 0.3355 | 0.6406 | 0.3594 | 0.5526 | 0.4474 | 0.7188 | 0.2813 |

#### Comparison with existing parcellations - external validation

In addition to the visual comparison, we compared the two solutions using external validation measures, taking the solution obtained from the second dataset as the ground truth (Table 5). Regardless of the measured selected, the similarity between the datasets was approximately 70%.

**Table 5.** Comparison of the parcellations for the first and second datasets in terms of the external validation measures (the parcellation for the second dataset served as the ground truth for the computation of the external validation measures for the first dataset).

| Comparison of solutions from the first and second datasets |  |  |  |  |  |  |  |  |
| --- | --- | --- | --- | --- | --- | --- | --- | --- |
| measure | Purity |  | Mutual Information |  | Jaccard |  | Fowlkes-Mallows |  |
| amygdala | left | right | left | right | left | right | left | right |
| value | 0.7713 | 0.8421 | 0.2676 | 0.3686 | 0.7713 | 0.8421 | 0.6668 | 0.7477 |

Moreover, we compared our parcellations with existing parcellations. As the masks used in the analysis differed in terms of the spatial distribution, the voxels that were localized in the amygdala in one mask could be assigned to another brain area (for example, in the comparison between the APMC mask and AT mask). For that reason, when we compared our results to those of existing parcellations, we needed to take into account this variability. We defined a limited comparison as one that compares only voxels that are included as parts of the amygdala in both masks, and a full comparison as one that takes into account all the voxels of the masks compared (in Figures 14 and 15, the regions are shaded in green and blue). We also provided a full comparison, taking into account all areas present in both masks (the whole areas in Figures 14 and 15).

**Fig. 14.**
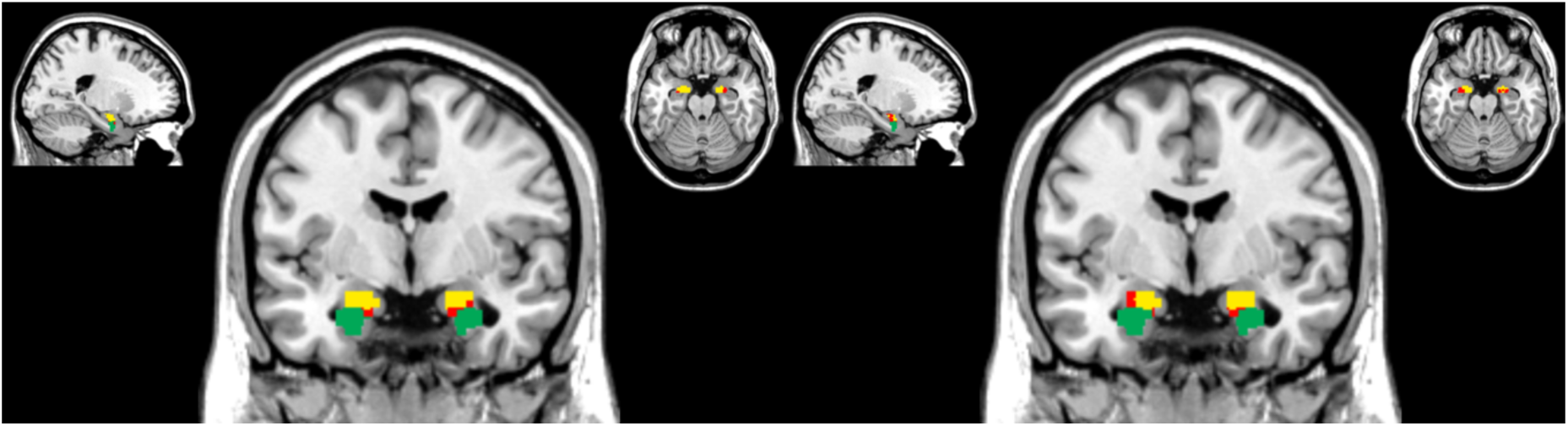
Comparison of the parcellations obtained for the first and second datasets with the parcellation obtained on the basis of structural connectivity patterns proposed by Dominic Bach and colleagues (2011). The green and yellow regions were classified in the same way in both parcellations. The red regions were classified as superficial (in our terminology a part of the DM) in the Bach et al. (2011) paper and VL (in the terminology of Bach et al.,2011 – deep) in our analysis. The results on the left were obtained on the basis of the first dataset, while those on the right were obtained on the basis of the second dataset. Coordinates: x = -21, y = -4, and z = -20; views: sagittal, coronal, and axial.

**Fig. 15.**
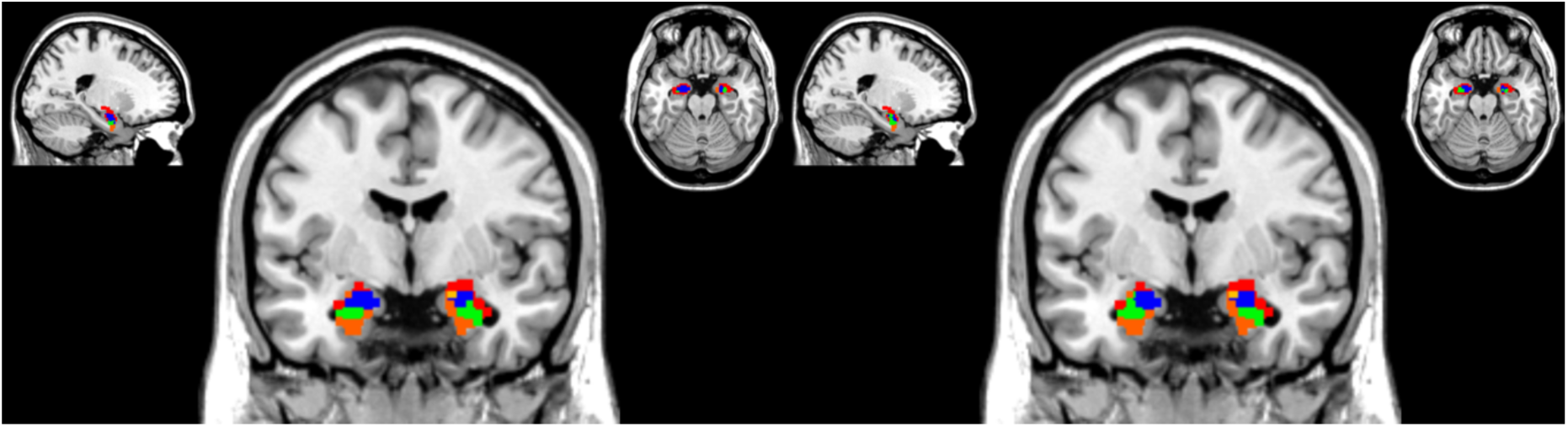
Comparison of the parcellations obtained for the first and second datasets with the parcellation obtained on the basis of cytoarchitecture by Katrin Amunts and colleagues (2005). The blue and green regions were classified in the same way in both parcellations; the rest of the colors denote that the solution was classified differently in the two datasets or that the voxel cannot be considered a part of the amygdala in one of the masks. Coordinates: x = -21, y = -4, and z = -20; views: sagittal, coronal, and axial.

The results of both the limited and full comparisons between the parcellation of the first dataset and the parcellation derived on the basis of structural connectivity patterns were the same and revealed a large degree of similarity between the two parcellations. The Fowlkes-Mallows score reached the level of approximately 75% with respect to the APMC mask, while the value of this indicator was much smaller than that obtained with respect to the SPM Anatomy toolbox mask. There are two reasons for this finding. First, the masks compared differ in terms of the spatial location of the voxels assigned to the amygdala – the mask from the Anatomy toolbox is significantly larger than the mask proposed by Bach et al. (2011). Second, parcellation on the basis of cytoarchitectonic features was performed on the three subparts, while our results indicated two functional subsystems within the amygdala. The values of the indices are presented in Table 6.

**Table 6.**
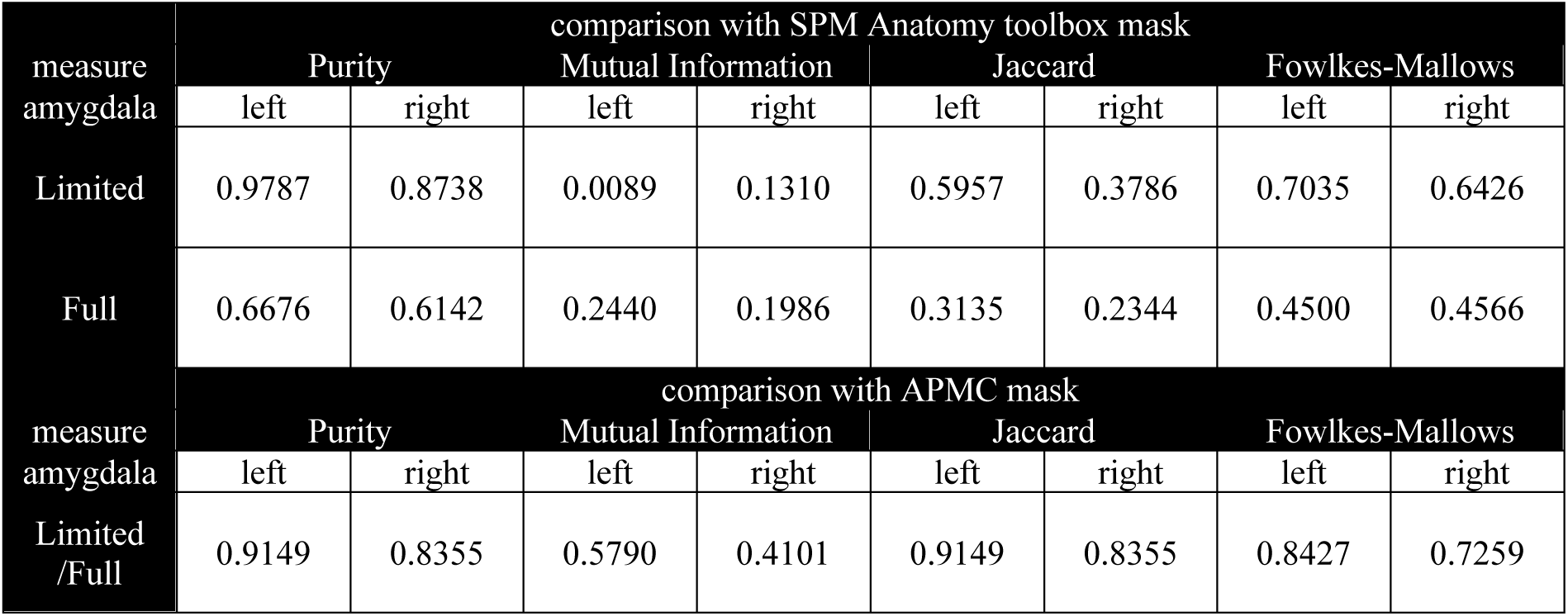
Values of the external validation measures of the amygdala parcellation for the first dataset.

We also calculated validation measures for the parcellation solution for the second dataset. As the proportions of sizes of the subdivisions were similar between the results of the second dataset and the SPM Anatomy toolbox, the external validation measures showed better results for this parcellations than for the comparison with the APMC parcellation (for example, value of Fowlkes-Mallows score for the limited comparison with the SPM Anatomy toolbox, Table 7). On the other hand, this comparison in the full scenario is similar to the results obtained on the first dataset. However, the Fowlkes-Mallows score was higher for the parcellation with first dataset than for the APMC parcellation and the parcellation with the second dataset. This finding proves that the results from the first dataset are more similar to the parcellation based on structural connectivity, while the results from the second dataset are more spatially congruent with the parcellation based on cytoarchitectonic features.

**Table 7.**
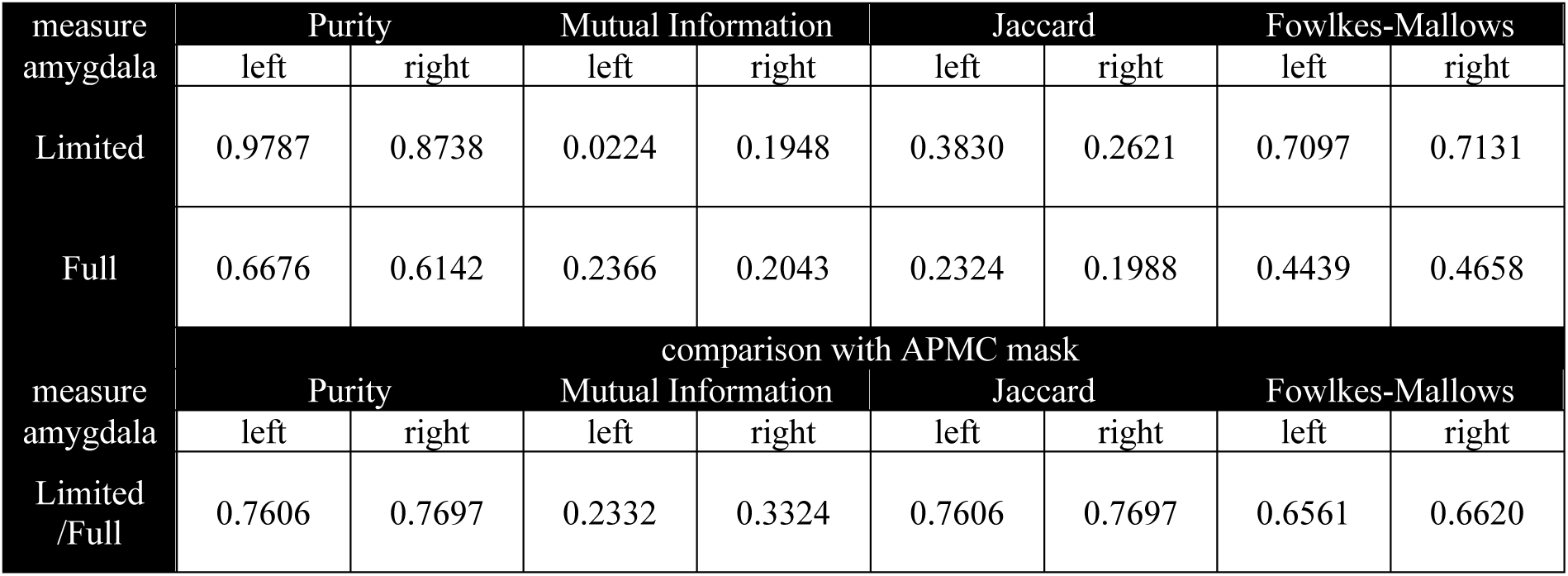
Values of the external validation measures of the amygdala parcellation for the second dataset.

#### Connectivity results of the third dataset

We showed that the obtained results are similar to those of previously existing parcellations. Until now, we labeled the obtained subdivisions as ventrolateral and dorsomedial subdivisions. To be able to align these names with anatomical nomenclature, we decided to perform a functional connectivity analysis on yet another dataset (the third dataset) with masks obtained in our analysis to identify the patterns of connections with different brain areas. We use the common areas between the ventrolateral and dorsomedial parts (as shown in Figure 12) and treated them as seeds in our analysis. To identify the distinct connectivity patterns characteristic to a specific amygdala subdivision, we compared all connections for the VL and DM subdivisions and determined which ones were had the strongest connections with a particular subpart.

The VL part of the amygdala in our data was characterized by functional connections with cortical regions – the frontal (middle frontal gyrus), parietal (supramarginal gyrus, cuneal cortex, precuneus, angular gyrus, superior parietal lobule), temporal (inferior temporal gyrus) and occipital regions (occipital fusiform cortex, supra- and intracalcarine cortex) (Table 8). Then, we repeated this analysis, but the main seed was in the mask of the dorsomedial amygdala and presented brain areas with significantly higher correlations with the spontaneous activity of this part than with the resting-state activity of the ventrolateral part. Thus, the dorsomedial part of the amygdala had stronger correlations with the subcortical structures such as the putamen, nucleus accumbens, thalamus and some cortical regions, such as the frontal medial cortex and postcentral gyrus. All of the correlations are presented in Table 9. In figure 16, the brain areas with stronger connections with the ventrolateral part are presented in red, and the brain areas with stronger connections with the dorsomedial subdivision are shown in blue.

**Table 8.**
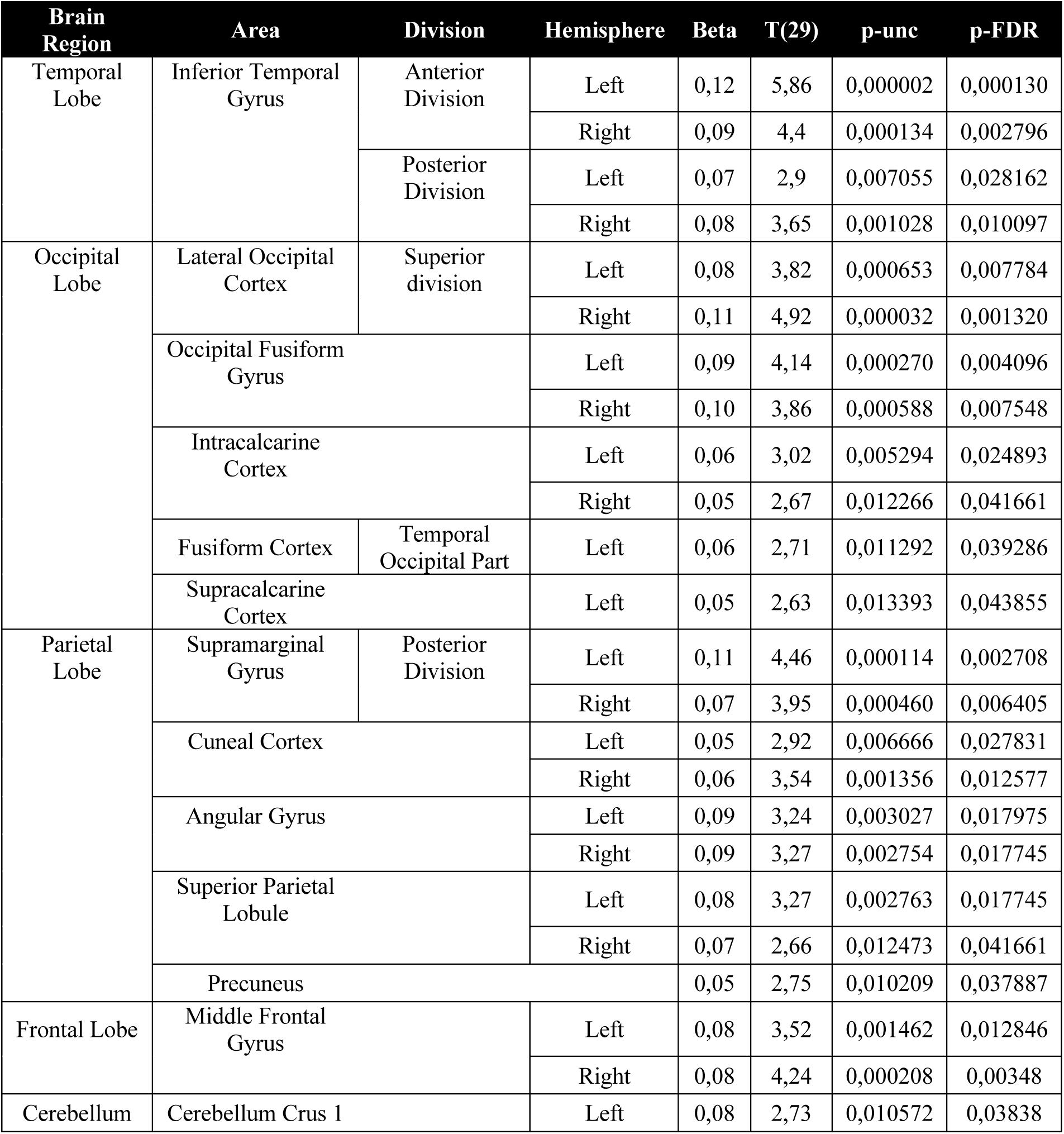
Functional connectivity of the ventrolateral part based on the common part between the datasets, which was applied to the third dataset.

**Table 9.** Functional connectivity of the dorsomedial part based on the common part between the two datasets, obtained for the third dataset.

| Brain Region | Area | Division | Hemisphere | Beta | T(29) | p-unc | p-FDR |
| --- | --- | --- | --- | --- | --- | --- | --- |
| Subcortical | Putamen |  | Left | 0,15 | 4,66 | 0,000065 | 0,001910 |
|  |  |  | Right | 0,15 | 4,64 | 0,000069 | 0,001910 |
|  | Hippocampus |  | Left | 0,13 | 3,70 | 0,000908 | 0,009478 |
|  |  |  | Right | 0,16 | 4,26 | 0,000197 | 0,003480 |
|  | Thalamus |  | Left | 0,08 | 3,13 | 0,003921 | 0,021125 |
|  |  |  | Right | 0,08 | 3,22 | 0,003121 | 0,017975 |
|  | Accumbens |  | Left | 0,08 | 3,17 | 0,003588 | 0,019975 |
|  |  |  | Right | 0,08 | 3,75 | 0,000792 | 0,008817 |
|  | Caudate |  | Left | 0,07 | 2,99 | 0,005669 | 0,025589 |
| Frontal Lobe | Frontal Medial Cortex |  |  | 0,08 | 3,48 | 0,001623 | 0,013551 |
|  | Central Opercular Cortex |  | Left | 0,07 | 3,04 | 0,005028 | 0,024694 |
|  |  |  | Right | 0,07 | 3,41 | 0,001951 | 0,015342 |
| Parietal Lobe | Parietal Operculum Cortex |  | Left | 0,07 | 3,32 | 0,002421 | 0,017578 |
| Temporal Lobe | Parahippocampal Gyrus | Posterior Division | Right | 0,11 | 3,26 | 0,002874 | 0,017775 |

**Fig. 16.**
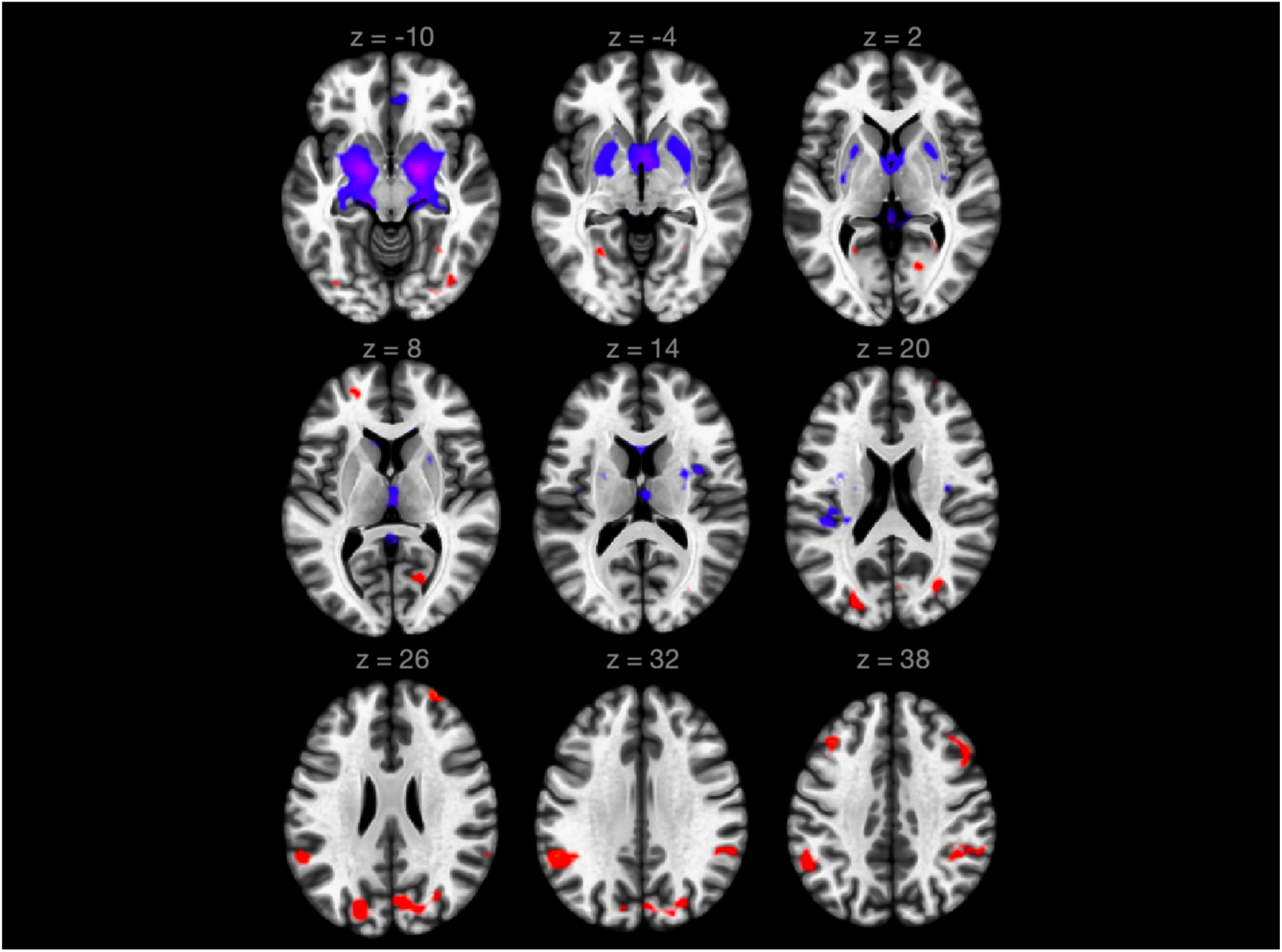
Visualization of the connections in the ventrolateral and dorsomedial parts for the third dataset.

## DISCUSSION

Our study introduced a novel approach to parcellate the human amygdala on the basis of the dynamic characteristics of the BOLD signals in the amygdala in voxels, as assessed by recurrence quantification analysis (RQA). We clearly partitioned the amygdala into two subdivisions: the ventrolateral (VL) and dorsomedial (DM) subdivisions. These names were given because of their location in the brain.

In our analysis, we assessed the system’s trajectory in the phase space and, subsequently, created a recurrence plot (Eckmann et al., 1987) for each voxel. Then, on the basis of the RPs, we computed several known measures for recurrence quantification analysis (RQA). From the set of these measures, Shannon’s entropy of the distribution of diagonal lines appears to be the most suitable for the differentiation of amygdala subdivisions on the basis of neuronal dynamics. It measures how unpredictable the trajectory of the analyzed system is. The larger the value of the entropy is, the more uniform the distribution of the lengths of the diagonal lines, and the more variable the lengths, which means that it is more difficult to predict how long the system will repeat a previously traveled sequence of states. Thus, Shannon’s entropy of diagonal lines focuses on the stability and regularity of a signal. For that reason, in our analysis, entropy was assessed for different features of data, whereas in previous investigations in the field of fMRI neuroimaging, the measure of entropy was directly applied to the BOLD time series (Wang et al., 2014; Mikoláš et al., 2012).

Regarding the masks we obtained, compared with the DM part, the VL part can be characterized by higher values of entropy. This finding may suggest that the VL subdivision is influenced by more brain systems. We tested this idea with the functional connectivity analysis, where masks of the VL and DM subdivisions were treated as seeds. VL activity was found to be correlated extensively with the cortical, occipital, parietal, temporal and frontal regions, whereas DM activity predicted activity primarily in the subcortical structures, such as the striatum, thalamus, and hippocampus. These two major amygdala complexes, with differential connectivity patterns, were consistent with connectivity patterns of the basolateral amygdala (BLA) and centromedial amygdala (CMA) (Roy et al., 2008). Anatomically recognized groups of amygdala nuclei (basolateral and centromedial) form networks for distinct functions via their unique pattern of interactions with other cortical and subcortical structures (LeDoux 2000; Davis & Whalen 2001). The BLA affectively evaluates sensory information and plays a critical role in the perception and regulation of emotionally salient stimuli via its projections to widely distributed cortical regions (LeDoux 2000; Davis & Whalen 2001; Jovanovic & Ressler, 2010).

In contrast, the CMA is essential for controlling the expression of emotional responses through projections to subcortical structures, including the thalamus, hypothalamus, striatum, and brainstem (LeDoux 2000; Davis & Whalen 2001). The fact that the VL seems to be similar to the BLA shows that the less stable dynamics (higher entropy of the length of repeated trajectories in the phase space) of the brain signal might be related to the contribution of evolutionarily younger brain subsystems associated with emotion expressions rather than the DM or the CMA.

The similarity in functional connectivity patterns between the VL and BLA, as well as between the DM and CMA, confirm the utility of our new parcellation approach. Moreover, our method yielded parcellations similar to previously described parcellations performed on the basis of structural connectivity patterns (Bach et al., 2011) and cytoarchitectonic characteristics (Amunts et al., 2005). It is a well-known fact that the spatial representation of brain areas highly correlates with but also partly differs from the results of structural and functional analysis (Honey et al., 2008). For that reason, the comparison of our results with those of parcellation on the basis of structural connectivity revealed high similarity but not identical solutions. Moreover, Bach and colleagues (2011) performed parcellation based on the structural connectivity patterns acquired via DWI and two regions with which connections differentiating the subdivisions of the amygdala were used. Nevertheless, the connectivity patterns of the masks obtained in our study correspond well with the brain circuits and networks of the BLA and CMA.

On the other hand, our results were less similar to those of parcellation based on cytoarchitectonic features than those of MRI-based research. This finding might be caused by the fact that the amygdala was partitioned into three parts in the work by Amunts et al. (2005), while we divided the amygdala into two subareas. Moreover, the mask generated with the SPM Anatomy toolbox was significantly larger than the masks we used for the purpose of our analysis. Thus, these two masks overlapped only partially, and these two parcellations were less similar to that in our results than was the APMC parcellation.

## CONCLUSIONS

In summary, we present a new parcellation approach for the human amygdala. We used RQA, which revealed the dynamic characteristics of the functional subsystems within the amygdala. This method also allowed us to further explore the nature of the BOLD signals in other brain areas. Although few related attempts have been made in the field of fMRI research (Bianciardi et al., 2007; Lombardi et al., 2017; beim Graben et al., 2019), this approach seems to be promising and useful for future research on functional brain organization among healthy subjects and patients with psychiatric disorders.

## CRediT authorship contribution statement

**Krzysztof Bielski:** Methodology, Investigation, Formal analysis, Visualization, Writing - original draft, Writing - review & editing, Funding acquisition.

**Sylwia Adamus:** Investigation, Formal analysis, Visualization, Writing - review & editing.

**Emilia Kolada:** Investigation, Writing - review & editing.

**Joanna Rączaszek-Leonardi:** Conceptualization, Methodology, Writing - review & editing, Supervision.

**Iwona Szatkowska:** Conceptualization, Writing - original draft, Writing - review & editing, Supervision, Project administration, Funding acquisition.

## Compliance with ethical standards

The study protocol was approved by the Ethic Committee for studies on humans of University of Social Sciences and Humanities (SWPS University) and University of Warsaw. Written informed consent was obtained from each subject before the study.

## Acknowledgments

We thank Marcel Falkiewicz for useful comments and support in data analysis. Additionally, we would like to thank Katarzyna Wisiecka and Pamela Sobczak for their help in acquiring the data, as well as Małgorzata Wordecha - Draps and the team from the Laboratory of Brain Imaging, Nencki Institute of Experimental Biology for support of team members.

