## Supplementary Materials for "Parcellation of the human amygdala using recurrence quantification analysis"

**APPENDICES**

**Recurrence Quantification Analysis (RQA) - description of the methods**

*Recurrence Quantification Analysis (RQA) – theoretical background.* The Recurrence quantification analysis (RQA) is a method derived from dynamic systems theory (DST), a kind of metatheory that is used to account for the properties of complex systems comprised of multiple nonlinearly interacting elements. RQA is one of the nonlinear techniques used for the "assessment or diagnosis of complex dynamical systems" (Zbilut & Webber, 1992). It is based on time series generated by a complex system and allows us to make inferences about the dynamical properties of that system. It has been extensively used to study heart rate dynamics (Zbilut, Thomason & Webber, 2002) in medical sciences and used in behavioral and cognitive research to identify differences in heart-rate activity (Konvalinka et al., 2011), EEG signals (Heunis et al., 2018) and even eye movement (Dale et al.,2011) and postural coordination (Shockley et al., 2009). Studies in which RQA was performed with MRI data are relatively rare (Bianciardi et al., 2007; Lombardi et al., 2017).

RQA focuses on describing dynamics of the system with quantitative measures of recurrence - when, how often and for how long an analyzed system returns to the previously visited states. RQA operates on a single time series (modifications of the technique have been developed for multiple time series, see Wallot, Roepstorff & Mønster, 2016), which is assumed to have been generated by a complex dynamical system. To compute measures of RQA, first, it is necessary to reconstruct the trajectory of the system in the phase space on the basis of a single time series with each point in the phase space representing one state of the system. The reconstruction is possible due to Taken’s theorem, which has proven that it is possible to reconstruct a higher-dimensional system consisting of multiple coupled variables from a single measured variable of that system by the time delay method (Zbilut & Webber, 1992). The next step is to visualize the reoccurrence of the system to similar values in this higher-dimensional space by constructing a so-called recurrence plot (RP). Thus, the RP presents the recurrent characteristics of the trajectory of a dynamical system in a multidimensional phase space on a 2D plane. RQA is a quantification of structures that appear in the RP. (An example of an RP graph is shown in Figure S1). RQA consists of several measures describing the density of the recurrence points, the proportion of diagonal and vertical lines in the RP, etc. (more details are provided in the subsection RQA measures).

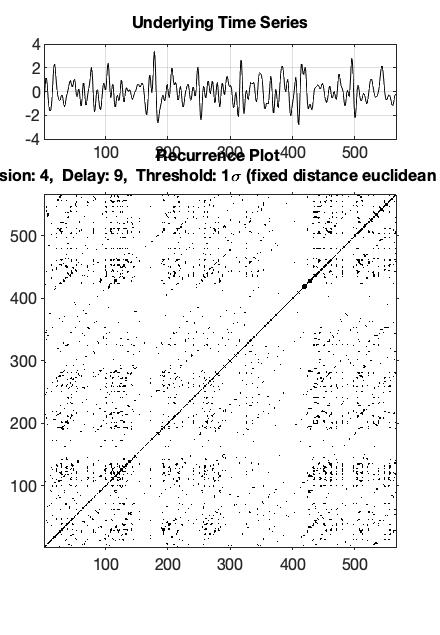

Fig. S1. A recurrence plot generated with the CRP toolbox (Marwan et al., 2007) and the signal of one of the voxels.

*RQA parameters estimation.* As stated above, the first step of RQA is reconstructing the trajectory of the system in the phase space. Based on Taken’s theorem mentioned above, we can reconstruct the trajectory of the whole system in a given multidimensional space by the method of delays (Takens, 1981). Thus, the next parameters we need to determine are how many time points we need to delay our time series and how many dimensions the system has to be reconstructed. To determine the former parameter, the average mutual information criterion (AMI) is used (Marwan et al., 2007). We compare how much information we lose by moving the time series by several time points. The point of the first local minimum is a desirable value of the time delay parameter (elbow point – Fig. S2). We computed this parameter using the rqatoolbox designed by Mike Richardson (Richardson et al., 2013). In our analysis, each time series had a different time delay value. The distribution of the time delay values for the different datasets is shown in the histogram (Fig. S3 – S6).

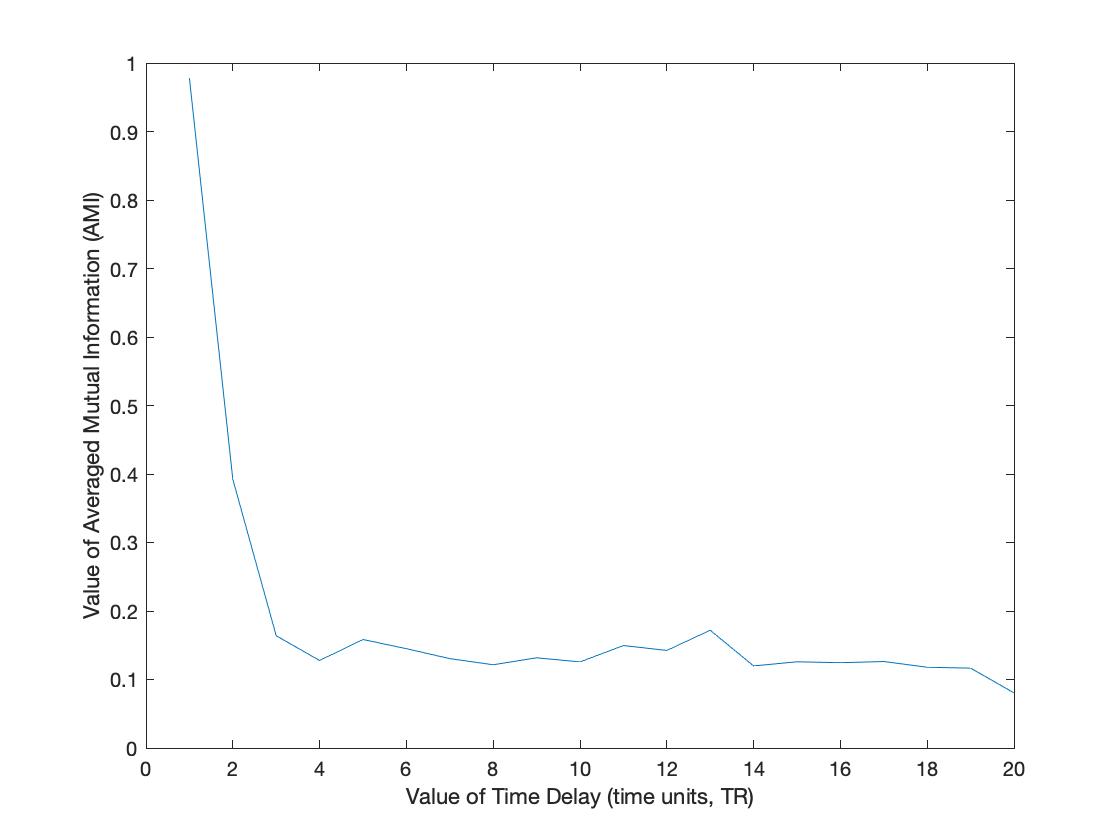

Fig. S2. Example of estimating the value of the time-delay parameter with a plot of the average mutual information criterion. The delay of the time points is shown on the x axis; the average mutual information loss is shown on the y axis.

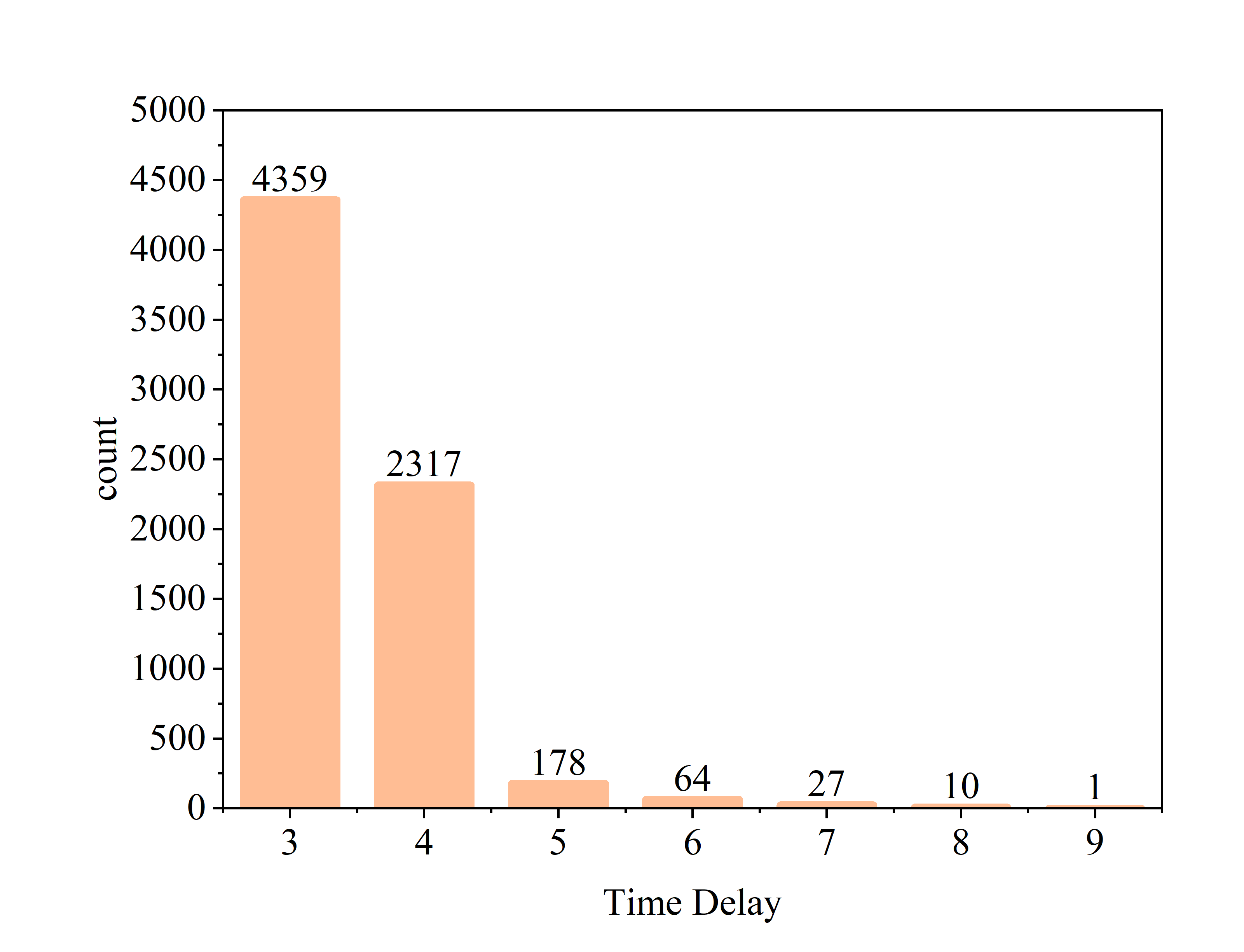

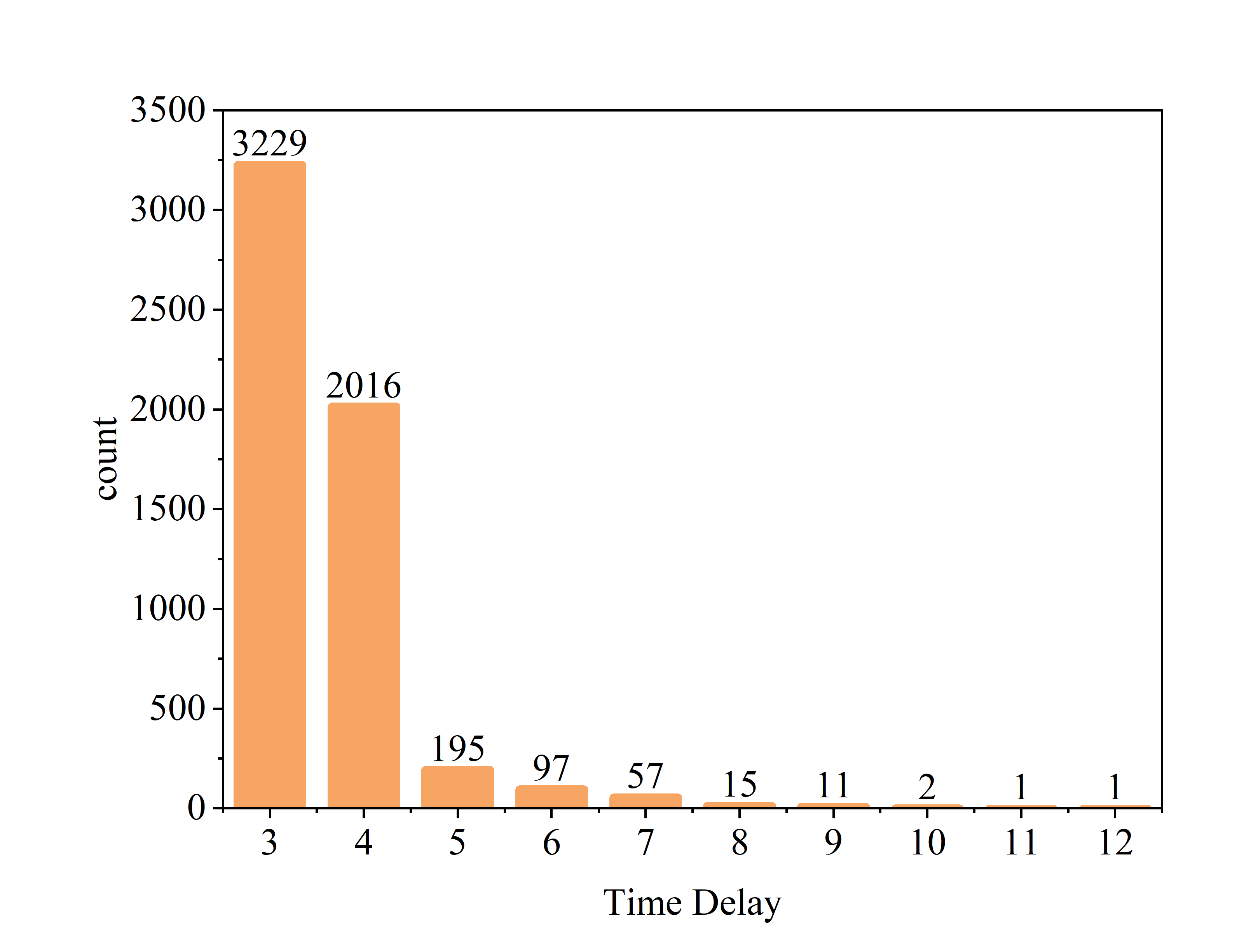

Fig. S3 and S4. Distribution of the time delay values for all voxels across subjects from the first dataset for the APMC mask for the left (light orange) and right (dark orange) amygdalae.

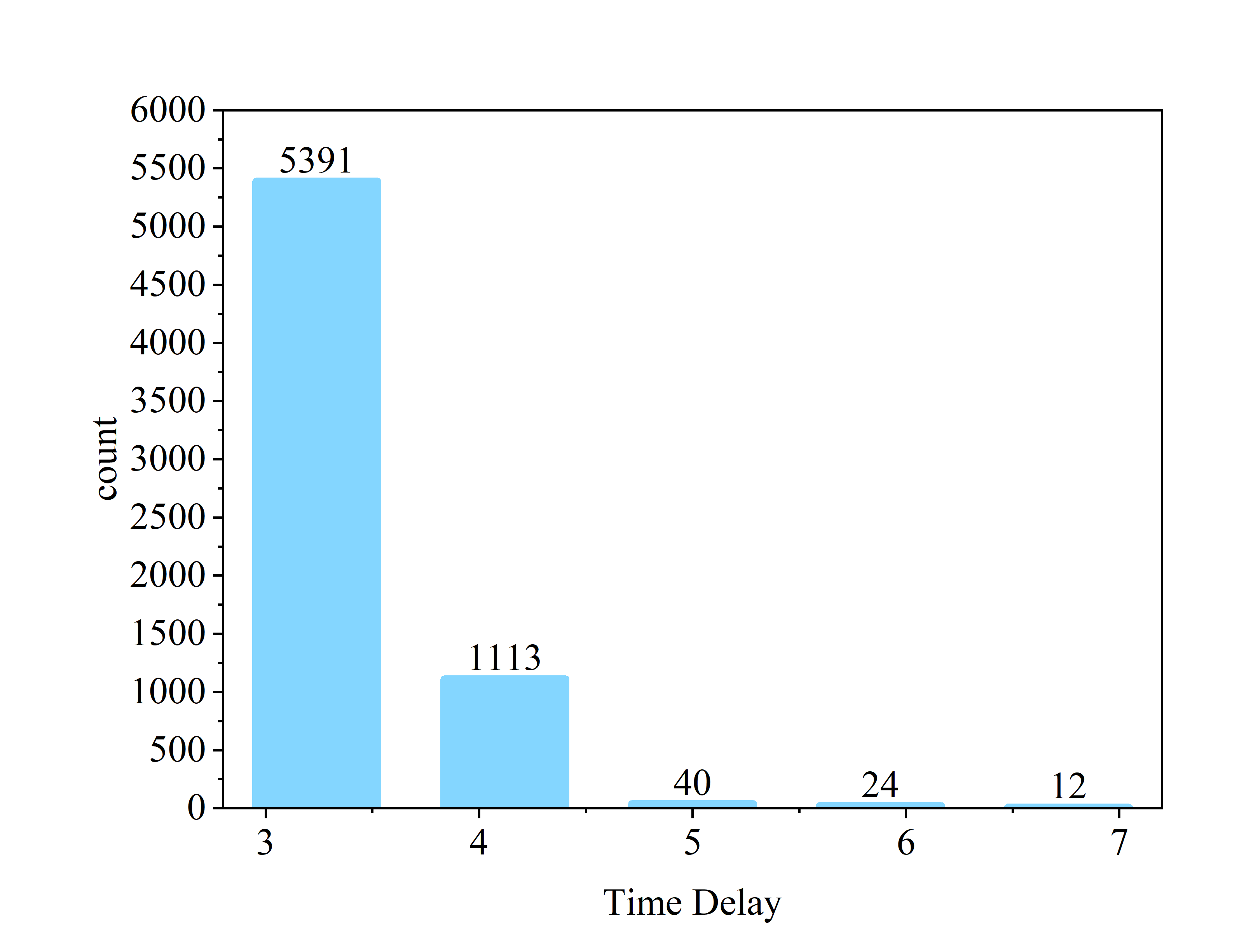

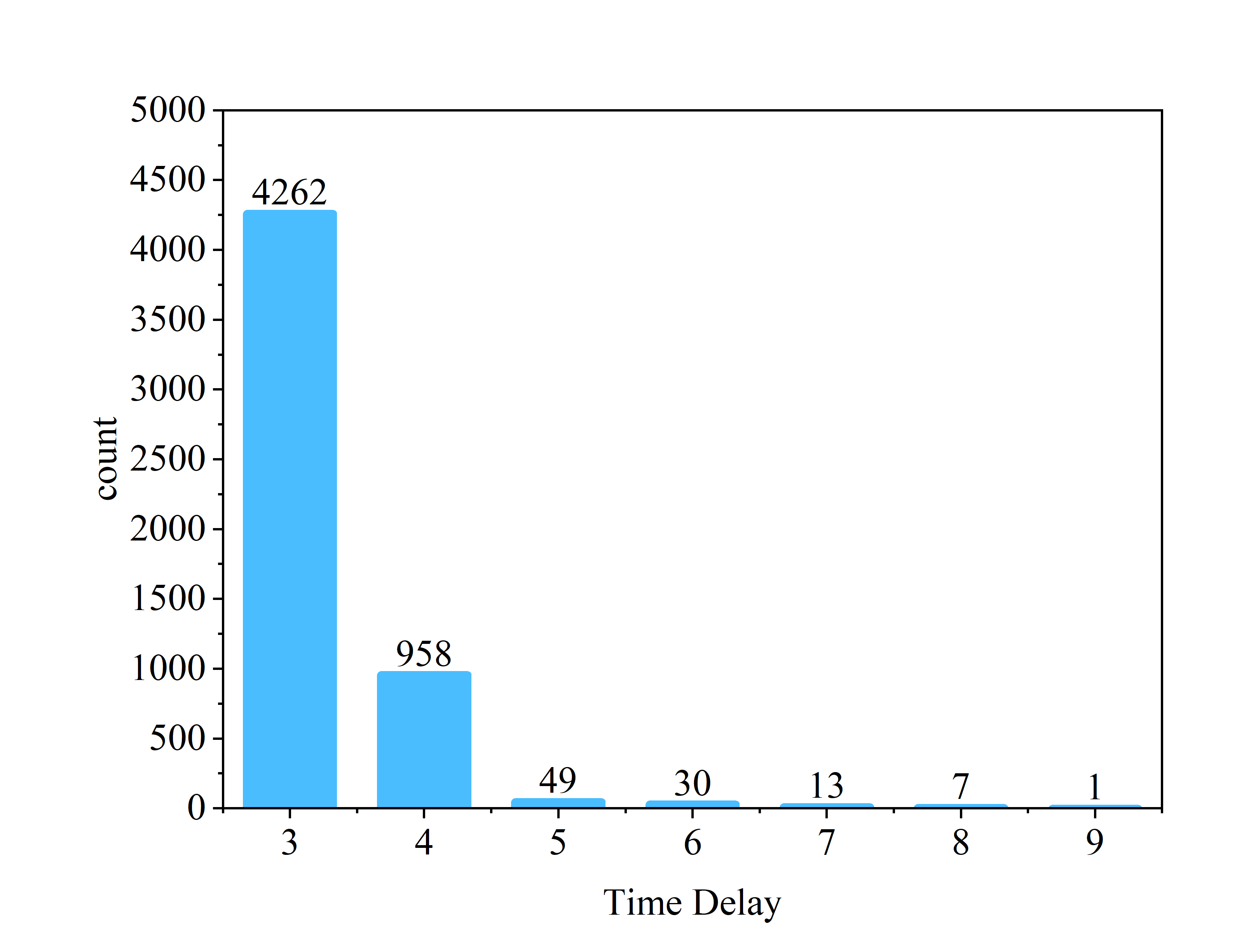

Fig. S5 and S6. Distribution of the time delay values for all voxels across subjects from the second dataset for the APMC mask for the left (light blue) and right (dark blue) amygdalae.

Next, we need to determine the dimensionality of the space in which the system’s trajectory will be reconstructed, e.g., estimate the embedding dimension. The most common method is that involving the false nearest neighbors’ criterion (Wallot & Monster, 2018). It follows the logic that two points in the original time series are treated as neighbors when they are located near each other. If they are transferred into the m-dimensional phase space (we delay the original time series m times with d time-points each, which corresponds to the delay), the distance between them might be substantially larger than in the lower dimensional representation of the time series. Increasing the embedding dimension parameter may lead to a larger distance between neighbors. However, if the neighbors embedded into the m-dimensional phase space cause a significant change in the distance between two points, then this pair of points is called a pair of false neighbors. Regarding the false nearest neighbors’ criterion, we focus only on the closest neighbors for the time points. When the number of dimensions increases, the number of false nearest neighbors decreases. In our analysis, we estimated the embedding dimensions for all voxels separately and then averaged them across the whole population. The distributions of embedding dimensions across voxels are provided in Tables S1 and S2.

| **value** | **left amygdala** | **right amygdala** |
| --- | --- | --- |
| **4** | 4981 | 3997 |
| **5** | 1972 | 1624 |
| **6** | 3 | 3 |

Table S1 Distribution of the embedding dimension values for all voxels across subjects from the first dataset.

| **value** | **left amygdala** | **right amygdala** |
| --- | --- | --- |
| **4** | 4620 | 3889 |
| **5** | 1958 | 1428 |
| **6** | 2 | 3 |

Table S2. Distribution of the embedding dimension values for all voxels across subjects from the second dataset.

The last parameter that needs to be estimated is the radius, which defines how large the area in which two time points are treated as being in the same state. There are several methods of obtaining the value of radius. First, “the rule of thumb” states that 10% of the maximum phase space diameter can be used as the value of radius (Zbilut & Webber, 1992). Some studies use a radius value equal to 20-40% of the standard deviation of the signal (Schinkel et al., 2008). Other works choose the value of the radius that returns the recurrence rate, allowing the structures to be identified on the RP. In our case, the radius was equal to or greater than 2% for each time series from a single amygdala (left/right) and a given mask. In Figures S7 and S8, the distributions of the recurrence rate in the whole analyzed population are shown. This method was also previously successfully applied to other types of data (Matassini et al.,2002; Marwan & Webber, 2015). Both the embedding dimension and radius were estimated using crptoolbox (MARWAN, N.: Cross Recurrence Plot toolbox for MATLAB®, version 5.22 (R32.2b), http://tocsy.pik-potsdam.de/CRPtoolbox/, accessed 2019-04-08.; Marwan et al., 2007).

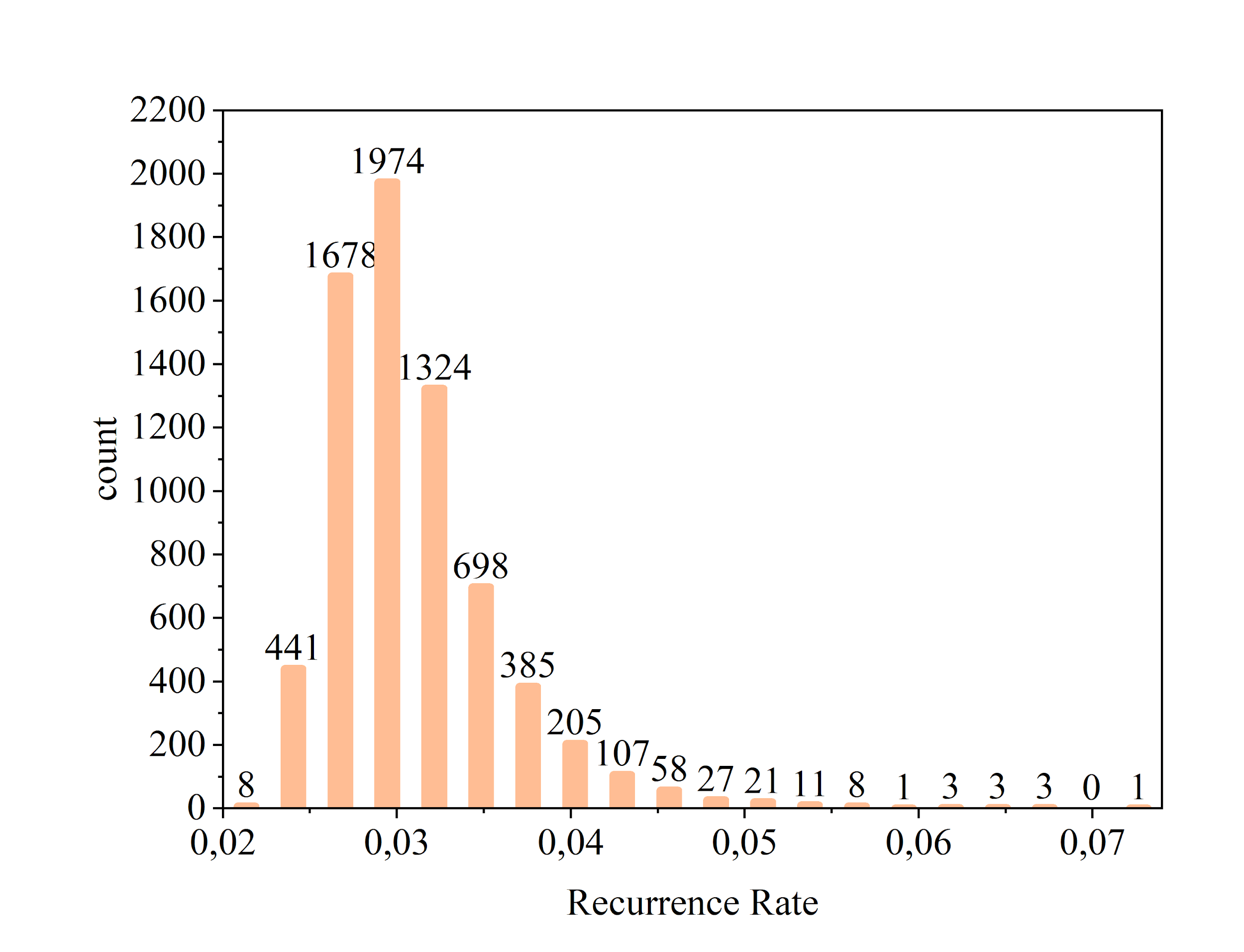

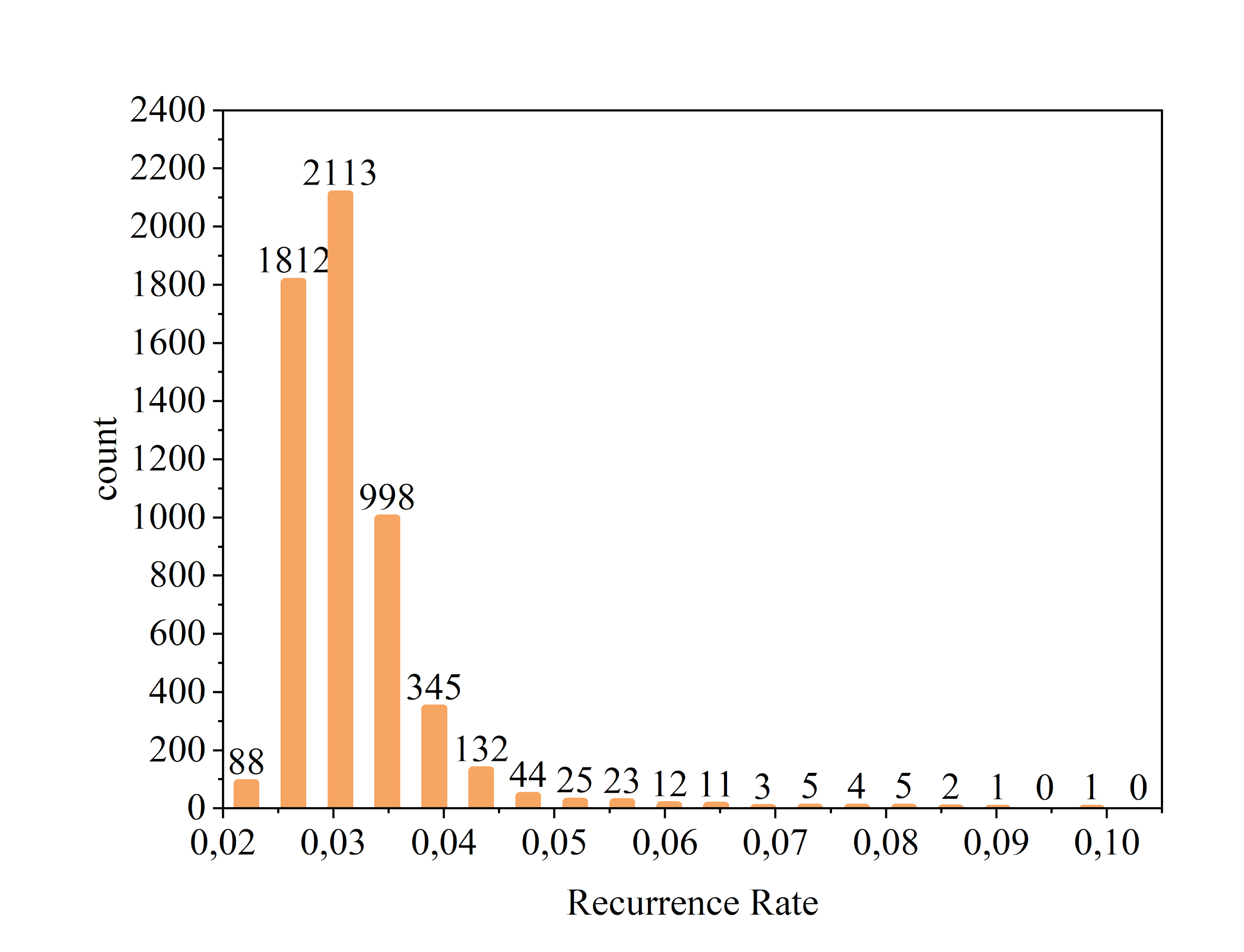

Fig. S7 and S8. Distribution of the recurrence rate values for all voxels across all subjects from the first dataset in the left and right amygdalae.

As our analysis was performed on the population level, our parameters needed to lead to comparable results between subjects. For that reason, the embedding dimension should have had the same value for each time series for all subjects for a given amygdala and mask. It is inappropriate to compare time series operating on the phase space with different numbers of dimensions. For that reason, similar to Marwan et al. (2007), we estimated the false nearest neighbors’ criterion and used the average to estimate the RQA measures for all subjects. Moreover, the value of the radius should have been the same in all voxels and all subjects. The values of the embedding dimension and radius used in the analysis are presented in Table S3.

|  | **left amygdala** | **right amygdala** |
| --- | --- | --- |
| **radius** | 1 | 1 |
| **embedding dimension** | 4 | 4 |

Table S3. Values of the radius and mean embedding dimension used to calculate the RQA measures for the first dataset.

*RQA measures.* After estimating the parameter values and performing RQA with these values, we created recurrence plots (RP), which show the timepoints that visit the same states and when (Figure S1). RQAs for each time series were performed using crptoolbox (Marwan et al., 2007). We computed the following measures:

- Recurrence rate (RR), which is a percentage of the recurrent points on the RP.

$$RR= \frac{1}{N^{2}}\sum_{i,j=1}^{N} R_{i,j}$$

Equation S1. Recurrence rate computed on the basis of RP. N – number of states in the RP; R is equal to 1 if there is the same value on the RP at time points i and j.

- %Determinism (%DET), which is the percentage of recurrent points that form diagonal lines (except the main diagonal - known as the line of identity, LOI). Deterministic systems can be characterized by repeated trajectories in the phase space. Many diagonal lines indicate that the system is deterministic, while short diagonal lines are characteristic of a randomly chaotic system.

$$\%DET= \frac{\sum_{l=l_{min}}^{N} lP(l)}{\sum_{l=1}^{N} lP(l)}$$

Equation S2. %DET measure: l is the length of a single diagonal line; N is number of diagonal lines of different lengths; P(l) is histogram of diagonal line with length equals to l.

- Average diagonal line length (AverageDiagonal), which is the average length of the diagonal lines, excluding the LOI, and shows how close trajectories are to each other and thus how predictive the time-series is.

$$L= \frac{\sum_{l=l_{min}}^{N} lP(l)}{\sum_{l=l_{min}}^{N} P(l)}$$

Equation S3. Average diagonal line length. The variable definitions are the same as those in Equation S2.

- Maximum diagonal line length (MaxLine), which is the maximum length of the diagonal line, excluding the LOI. It demonstrates how divergent the system; the longer the line is, the smaller the divergence.
- Shannon's entropy of the frequency distribution of the diagonal line lengths (ENTR), which is derived from information theory and reflects the level of complexity of the deterministic structure of the system, as expressed by the diagonal lines in the recurrence plot (RP). The higher the value is, the more chaotic and unpredictable the system.

$$ENTR= \sum_{l=l_{min}}^{N} p(l)\log p(l)$$

Equation S4. Shannon’s entropy of diagonal lines. The variable definitions are the same as those in Equation S2.

- Laminarity (LAM), which is the percentage of recurrent points that form vertical lines. This measure indicates how stationary the system is over time as a percentage. The higher the LAM value, the more often the system remains in the same states.

$$LAM= \frac{\sum_{v=v_{min}}^{N} vP(v)}{\sum_{v=1}^{N} vP(v)}$$

Equation S5. Laminarity: v is the length of a single vertical line; N is number of vertical lines of different lengths; P(l) is histogram of vertical line with length equals to l.

- Trapping time (TT), which is the average length of the recurrent points that form vertical lines. Analogous to the average diagonal line length, it states how long the system on average is in one state.

$$TT= \frac{\sum_{v=v_{min}}^{N} vP(v)}{\sum_{v=v_{min}}^{N} P(v)}$$

Equation S6. Trapping time. The variable definitions are the same as those in Equation S5.

**Internal Validity Measures**

To evaluate how good clustering classification is, it is necessary to compute whether points within one cluster are characterized by relatively small distances and distances between points in different clusters are as high as possible. It can be presented using measures such as BetaCV, normalized cut and the silhouette coefficient.

- BetaCV is the ratio of the average distance between points within one cluster and average distance between points in different clusters. The smaller the value is, the better the clustering.

$$BetaCV= \frac{W_{in}/N_{in}}{W_{out}/N_{out}}$$

Equation S7. Value of the BetaCV indicator. The W and N values are computed in Equations S8-S11.

$W_{in}= \frac{1}{2}\sum_{i=1}^{k} W(C_{i},C_{i}$)

Equation S8. Value of the sum of intracluster weights over all clusters. W is the sum of weights between points within a cluster; k – number of clusters

$W_{out}= \frac{1}{2}\sum_{i=1}^{k} W(C_{i},\bar{C}_{i}$)

Equation S9. The sum of the intercluster weights across all clusters. W is the sum of weights between points between two clusters; The line above C indicates C values other than Ci; k – number of clusters

$$N_{in}=\sum_{i-1}^{k} \left( \begin{matrix} n_{i} \\ 2 \end{matrix} \right)$$

Equation S9. The number of distinct intracluster weights, where n is the number of weights in a given cluster and k is the number of clusters.

$$N_{out}=\sum_{i=1}^{k-1} \sum_{j=i+1}^{k} n_{i}n_{j}$$

Equation S10. Value of the number of distinct intercluster weights, where n is the number of weights in a given cluster and k is the number of clusters.

- Normalized cut is the ratio of the sum of intracluster distances and the sum of inter- and intracluster distances. It takes values from 0 to 1. The larger the value is, the better.

$$NormalizedCut= \sum_{i=1}^{k} \frac{1}{\frac{W(C_{i},C_{i})}{W(C_{i},\bar{C}_{i})}+1}$$

Equation S11. The normalized cut formula, were W is a sum of weights between points in a given cluster C_i_.

- The silhouette coefficient is the most restrictive criterion, taking into account the mean intracluster distances and mean nearest-cluster distance, and it is computed following this equation: (b-a)/max(a,b). Thus, good clustering does not occur when the distance within one cluster exhibit large variance. It takes a value between -1 and 1, where 0 is a completely random result, 1 indicates the best level of clustering, and -1 indicates that the points were assigned to bad labels.

$$Silhouette Coefficient= \frac{(b-a)}{\max(a,b)}$$

Equation S12. The silhouette score, where a is the mean intracluster distance and b is the mean nearest-cluster distance.

All measures were computed using Python 3.7.

| **Ranking** | **Measure** | **BetaCV** |
| --- | --- | --- |
| 1 | ENTR | 0.615287 |
| 2 | LAM | 0.650641 |
| 3 | TT | 0.686738 |
| 4 | AverageDiagonal | 0.696475 |
| 5 | DET | 0.706781 |
| 6 | MaxLine | 0.711334 |

Table S3. The values of BetaCV across solutions in the left amygdala for the first dataset.

| **Ranking** | **Measure** | **BetaCV** |
| --- | --- | --- |
| 1 | ENTR | 0.605200 |
| 2 | DET | 0.619672 |
| 3 | TT | 0.628541 |
| 4 | AverageDiagonal | 0.629797 |
| 5 | LAM | 0.777349 |
| 6 | MaxLine | 0.784097 |

Table S4. The values of BetaCV across solutions in the right amygdala for the first dataset.

| **Ranking** | **Measure** | **NormalizedCut** |
| --- | --- | --- |
| 1 | ENTR | 0.766615 |
| 2 | LAM | 0.747182 |
| 3 | DET | 0.737788 |
| 4 | TT | 0.734212 |
| 5 | MaxLine | 0.723700 |
| 6 | AverageDiagonal | 0.714626 |

Table S5. The values of NormalizedCut across solutions in the left amygdala for the first dataset.

| **Ranking** | **Measure** | **NormalizedCut** |
| --- | --- | --- |
| 1 | ENTR | 0.766903 |
| 2 | AverageDiagonal | 0.762858 |
| 3 | DET | 0.762825 |
| 4 | TT | 0.750250 |
| 5 | LAM | 0.688450 |
| 6 | MaxLine | 0.634786 |

Table S6. The values of NormalizedCut across solutions in the right amygdala for the first dataset.

| **Ranking** | **Measure** | **Silhouette** |
| --- | --- | --- |
| 1 | ENTR | 0.3554531 |
| 2 | LAM | 0.3257752 |
| 3 | TT | 0.2998493 |
| 4 | AverageDiagonal | 0.2838704 |
| 5 | DET | 0.2588635 |
| 6 | MaxLine | 0.257230 |

Table S7. The values of the silhouette coefficient across solutions in the left amygdala for the first dataset.

| **Ranking** | **Measure** | **Silhouette** |
| --- | --- | --- |
| 1 | ENTR | 0.359694 |
| 2 | TT | 0.348161 |
| 3 | DET | 0.340442 |
| 4 | AverageDiagonal | 0.331719 |
| 5 | LAM | 0.222151 |
| 6 | MaxLine | 0.208149 |

Table S8. The values of the silhouette coefficient across solutions in the right amygdala for the first dataset.

**External Validation Measures**

We compare different parcellations using four metrics of external validation. The external validation relates to the distinction between the obtained solution by clustering and the existing division of points into clusters labeled the ground truth solution. The higher the value of the external validation measures, the more similar to the ground truth the given clustering is. However, the similarity between the obtained solution and ground truth can be computed by taking into account different aspects of this comparison. Here, we present the external validation measures that we computed in our analysis:

1. Purity, which is the mean of the proportion of common parts between two parcellations (the parcellation obtained by us and the ground truth parcellation). To compute purity, the proportions of points from the ground truth cluster within the borders of the cluster need to be determined in the analyzed solution. We took into account only the ground truth cluster, which had the largest number of points (Equation S13). The global value of purity was computed by averaging the weighted purity values across all clusters (Equation S14).

$${purity}_{i}=\frac{1}{n_{i}}\max_{j=1}\left\{ n_{ij} \right\}$$

Equation S13. Purity of a single cluster i. n_i_ – number of points within the cluster i; n_ij_ – number of points in the cluster i which comes from the ground truth cluster j.

$$purity=\sum_{i=1}^{r} \frac{n_{i}}{n}{purity}_{i}$$

Equation S14. Purity of all clusters. n_i_ – number of points within cluster i; n – number of all points; r – number of all clusters

2. Mutual information (MutualInfo), which reflects the shared information between the clustering results and the ground truth solution. It quantifies the relation between the value of joint probability of the clustering results and ground truth and the expected probability joint probability of the clustering results and ground truth when we assume independence between them.

$$MutualInfo\left( C,T \right)=\sum_{i=1}^{r} \sum_{j=1}^{k} p_{ij}log\left( \frac{p_{ij}}{p_{C_{i}}*p_{T_{j}}} \right)$$

Equation S15. Mutual information of clustering; p_ij_ – observed joint probability; p_Ci_, p_Tj_ – expected joint probability with assumption that they are independent.

3. The Jaccard coefficient (Jaccard), which reflects the fraction of true positive point pairs and does not take into account true negatives. In other words, it is a coefficient that is associated with the relative number of points that are labeled the same way in the clustering results and ground truth (Equation S16).

$$Jaccard=\frac{TP}{\left( TP+FN+FP \right)}$$

Equation S16. The Jaccard coefficient. TP – True positives – two points classified in the same cluster both in the clustering results and ground truth, FN – false negatives – two points classified as belonging to the same ground truth cluster but not in the case of clustering; FP – false positives – two points classified as not belonging to the same ground truth cluster but in the case of clustering, they are located within the borders of the same cluster.

4. Fowlkes-Mallows score, which takes into account the false and true positives/negatives.

$$Fowlkes-Mallows=\frac{TP}{\sqrt{\left( TP+FN \right)\left( TP+FP \right)}}$$

Equation S17. Fowlkes-Mallows score. The variable definitions are provided in the explanation for Equation S10.
